# Preserved motor representations after paralysis

**DOI:** 10.1101/2021.10.07.463105

**Authors:** Charles Guan, Tyson Aflalo, Carey Y. Zhang, Emily R. Rosario, Nader Pouratian, Richard A. Andersen

## Abstract

Neural plasticity allows us to learn skills and incorporate new experiences. What happens when our lived experiences fundamentally change, such as after a severe injury? To address this question, we analyzed intracortical population activity in a tetraplegic adult as she controlled a virtual hand through a brain-computer interface (BCI). By attempting to move her fingers, she could accurately drive the corresponding virtual fingers. Neural activity during finger movements exhibited robust representational structure and dynamics that matched the representational structure, previously identified in able-bodied individuals. The finger representational structure was consistent during extended use, even though the structure contributed to BCI decoding errors. Our results suggest that motor representations are remarkably stable, even after complete paralysis. BCIs re-engage these preserved representations to restore lost motor functions.

## Introduction

A central question in neuroscience is how experience affects the nervous system. Studies of this phenomenon, plasticity, were pioneered by Hubel and Wiesel, who found that temporary visual occlusion in kittens can induce lifelong reorganization of the visual cortex^1^. Their results demonstrated that the developing brain, rather than being genetically preprogrammed, is surprisingly malleable to external inputs.

Subsequent studies showed that other brain regions are also plastic during early development, but it is less clear how plastic the nervous system remains into adulthood. Visual occlusion in adult cats does not reorganize the visual cortex, and lesion studies of the adult visual cortex have arrived at competing conclusions of reorganization and stability^2–5^. A similar discussion continues regarding the primary somatosensory cortex (S1). Amputation was classically thought to modify the topography of body parts in S1, with intact body parts taking over cortical areas originally dedicated to the amputated part^6–8^. However, recent human neuroimaging studies have challenged the extent of this remapping, arguing that sensory topographies in S1 largely persist even after complete sensory loss^9^. Thus, the level of plasticity in the adult nervous system is still an ongoing investigation.

Understanding plasticity is further necessary to develop brain-computer interfaces (BCIs) that can restore sensorimotor function to paralyzed individuals^10^. First, paralysis disrupts movement and blocks somatosensory inputs to motor areas, which could cause neural reorganization^8^. Second, BCIs create direct motor pathways that bypass supporting cortical, subcortical, and spinal circuits, fundamentally altering how the cortex affects movement. These changes raise an important question: do paralyzed BCI users need to learn a fundamentally new skillset^11^, or can they leverage their pre-injury motor repertoire^12^? If paralyzed BCI users can still engage natural movement representations, then adult motor representations may be more stable than classical amputation studies^6^ suggest. Assistive BCIs should then leverage the natural structure of these stable representations^12^.

Here, we test whether the neural representational structure of BCI finger movements in a tetraplegic individual matches that of able-bodied individuals during overt movements or whether it follows the optimal representational structure defined by the BCI task^13^. In able-bodied individuals, the cortical representational structure of finger movements follows the natural statistics of movements^14,15^. Within a BCI environment, the experimenter defines the movement statistics, which can be independent of biomechanics or before-injury motifs. We report that the neural representational structure of BCI finger movements in a tetraplegic individual matches that of able-bodied individuals. This match was stable across sessions, even though the measured representational structure contributed to errors in the BCI task. Furthermore, the neural representational dynamics matched the temporal profile expected in able-bodied individuals. Our results reveal that adult motor representations are preserved even after years without use.

## Results

### Intracortical recordings during finger flexion

We recorded single and multi-neuron activity (95.8 +/- s.d. 6.7 neurons per session over 10 sessions) from participant X (abbreviation: PX) while she attempted to move individual fingers of the right hand. We recorded from a microelectrode array implanted in the left (contralateral) posterior parietal cortex (PPC) at the junction of the postcentral and intraparietal sulci (PC-IP, Supplementary Figure 1), a region thought to specialize in the planning of grasping movements^16–19^.

Each recording session started with an initial calibration task (Supplementary Figure 2, Methods). On each trial, we used a computer screen to present a text cue (e.g., “T” for thumb), and the participant immediately attempted to flex the corresponding finger, as though pressing a key on a keyboard. Because PX previously suffered a C3-C4 spinal cord injury resulting in tetraplegia, her movement attempts did not generate overt motion. Instead, PX attempted to move her fingers as though they were not paralyzed.

These attempted movements resulted in distinct neural activity patterns across the electrode array. To enable BCI control, we then trained a linear classifier (Methods) to identify the attempted finger movement from the neural firing rates. The participant then performed several rounds of a similar attempted finger flexion task, except that 1) the trained classifier now provided text feedback of its predicted finger and 2) the task randomized the visual cue location (Figure 1a and Methods). We repeated this online-control finger flexion task over multiple sessions (408 +/- s.d. 40.8 trials/session over 10 sessions) and used this data for our offline analyses. PX also performed a control task, identical in structure except that PX attended to cues without performing the instructed movements.

**Figure 1.**
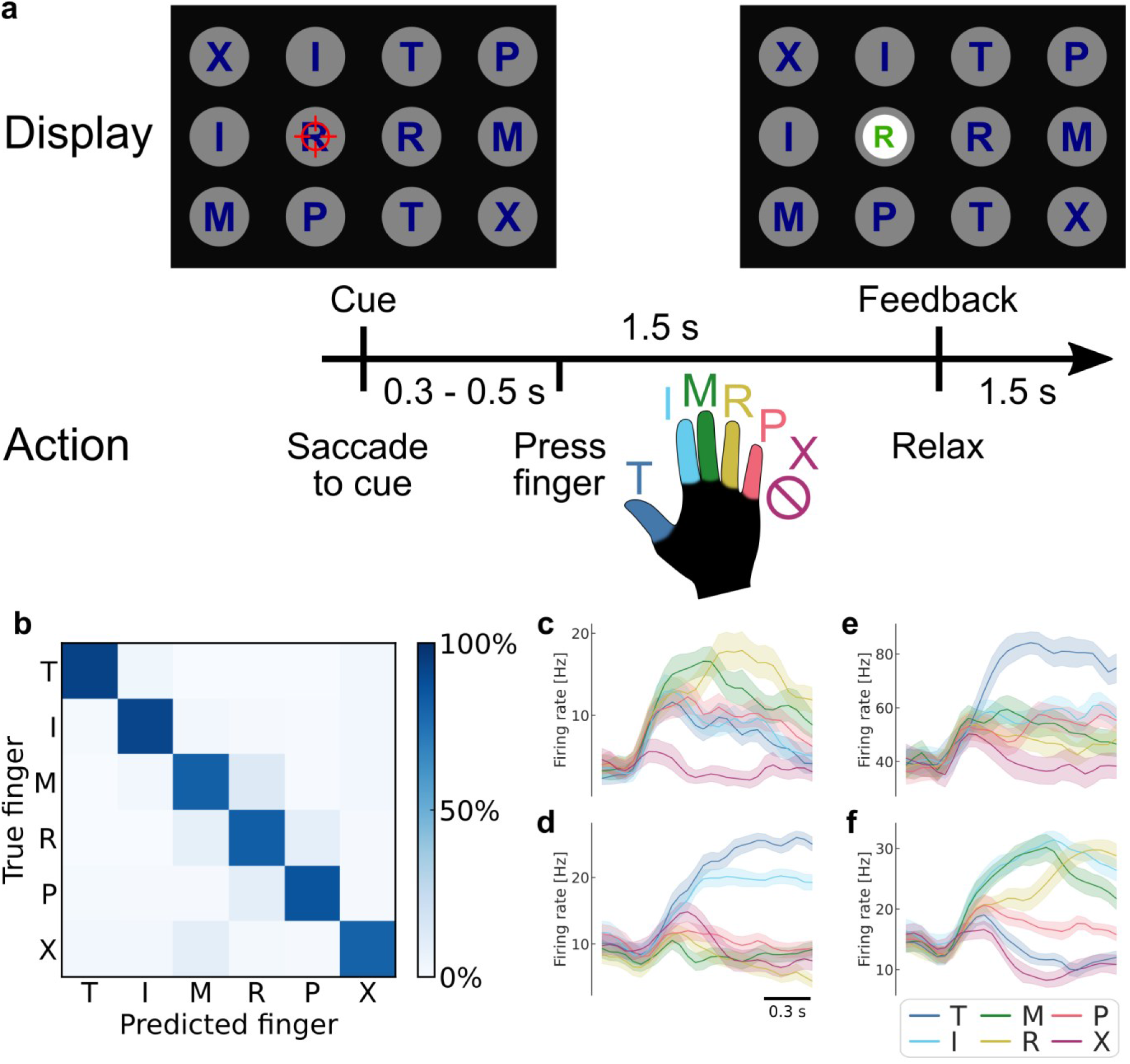
Robust brain-computer interface (BCI) control of individual fingers. (**a**) Main finger flexion task. When a letter was cued by the red crosshair, the participant looked at the cue and immediately attempted to flex the corresponding digit of the right (contralateral) hand. We included a null condition “X,” during which the participant looked at the target but did not move her fingers. Visual feedback indicated the decoded finger 1.5 seconds after cue presentation. To randomize the saccade location, cues were located on a grid (3 rows, 4 columns) in a pseudorandom order. The red crosshair was jittered to minimize visual occlusion. (**b**) Confusion matrix showing robust in-session BCI finger control (86% overall accuracy, 4016 trials aggregated over 10 sessions). Each entry (*i, j*) in the matrix corresponds to the ratio of movement *i* trials that were classified as movement *j*. (**c-f**) Mean firing rates for 4 example neurons, color-coded by attempted finger movement. Shaded areas indicate 95% confidence intervals (across trials of one session). Gaussian smoothing kernel (50-ms SD).

### Accurately decoding fingers from PPC single-neuron activity

High accuracy during online control (86% +/- s.d. 4% over 10 sessions; chance = 17%) (Figure 1b) and in offline cross-validated classification (89% +/- s.d. 2%) (Supplementary Figure 3) demonstrated that the finger representations were reliable and linearly separable. During the calibration task, cross-validated classification was similarly robust (accuracy = 95% +/- s.d. 3%; chance = 20%) (Supplementary Figure 3). These finger representations were robust across contexts and could be used in a range of environments, including to move a virtual reality avatar hand (Supplementary Figure 4).

At the single-neuron level, most (89%) neurons were significantly tuned to individual finger flexion movements (significance threshold: P < 0.05, FDR corrected) (Supplementary Figure 5). The example neurons in Figure 1c-f show that neurons could be tuned to one or more fingers and that tuning profiles could change in time.

To confirm that the observed neural responses could not be explained by visual confounds, we verified that we could not discriminate between fingers during the control task (Supplementary Figure 6). Furthermore, we could not decode the gaze location during the finger classification time window in the standard online-control task (Supplementary Figure 6). Thus, reliable finger representations emerged from the participant’s movement attempts.

### Finger representational structure matches the structure of able-bodied individuals

Having discovered that PC-IP neurons represent finger movements, we next investigated how these neural representations were functionally organized and how this structure relates to pre-injury movement.

Here, we turn to the framework of representational similarity analysis (RSA)^20,21^. RSA quantifies neural representational structure by the pairwise distances between each finger’s neural activity patterns (Figure 2a). These pairwise distances form the representational dissimilarity matrix (RDM), a summary of the representational structure. Importantly, these distances are independent of the original feature types (for example, electrode or voxel measurements), allowing us to compare representational structures across subjects and across recording modalities^22^.

**Figure 2.**
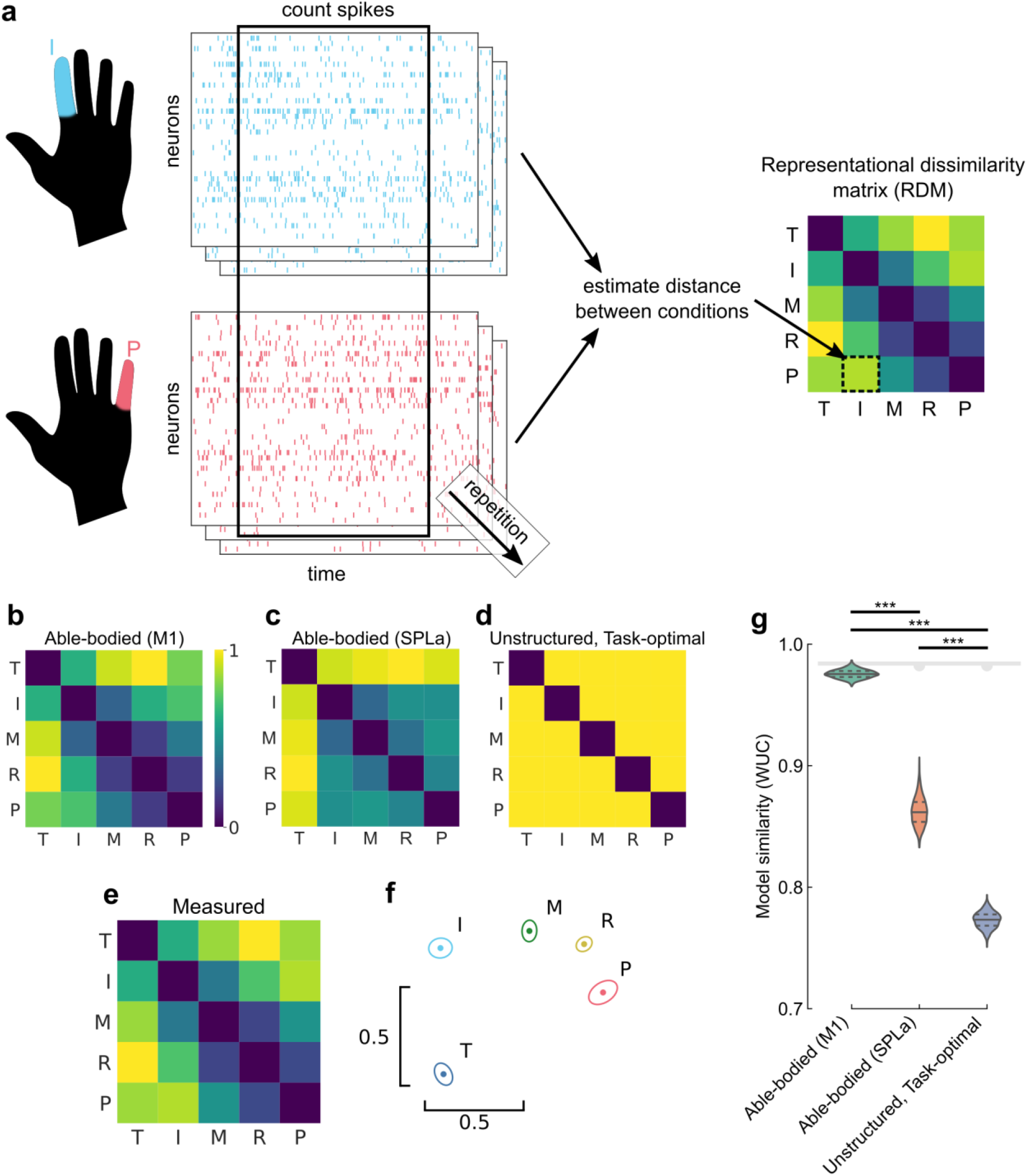
Representational structure during BCI finger control matches the structure of able-bodied individuals. (**a**) To construct the representational dissimilarity matrix (RDM), a vector of firing rates was constructed for each trial. Repetitions were collected for each condition. Then, pairwise distances were estimated between conditions using a cross-validated dissimilarity metric. This process was repeated to generate an RDM for each session. We drop the No-Go condition (X) here to match previous finger studies^14,23^. (**b**) Representational structure hypothesized by the preserved-representation hypothesis: average RDM for a finger-press task using 7T fMRI in 7 able-bodied individuals^14^. Max-scaled to [0, 1]. (**c**) Another possible representational structure hypothesized by the preserved-representation hypothesis: average RDM for a finger-press task using 3T fMRI in 20 able-bodied individuals^28^. Max-scaled to [0, 1]. (**d**) Representational structure hypothesized by the de-specialization and task-optimal hypotheses: pairwise-equidistant RDM. Max-scaled to [0, 1]. (**e**) Representational structure measured in our experiment: cross-validated Mahalanobis distances (Methods) between neural activity patterns in a tetraplegic individual, averaged across 10 recording sessions. Max-scaled to [0, 1]. (**f**) Intuitive visualization of the distances in (e) using multidimensional scaling (MDS). Ellipses show mean +/- s.d. (10 sessions) after Generalized Procrustes alignment (with scaling) across sessions. (**g**) Measured RDMs (d) match the able-bodied BOLD RDM (b) better than they match the unstructured (null) model (c), as measured by the whitened unbiased cosine similarity^27^ (WUC) (Methods). Difference was significant (P = 1.8 × 10^-10^, two-tailed t-test, 1000 bootstrap samples over 10 sessions). Violin plot: solid horizontal lines indicate the mean WUC over bootstrap samples, and dotted lines indicate the first and third quartiles. Noise ceiling: Gray region estimates the best possible model fit (Methods). Downward semicircle indicates that the unstructured model fit is significantly lower than the noise ceiling (two-tailed t-test, P < 0.001, Bonferroni-corrected for 3 model comparisons). Y-axis: even when the model and data RDMs are uncorrelated, WUC can be close to one^27^. For convenience, a similar figure using a correlation-based similarity metric is shown in Supplementary Figure 10.

We use RSA to test between four hypotheses: 1) we predicted that the finger representation would match the characteristic topographic organization identified in able-bodied individuals^14^ (Figure 2b) that follows the natural statistics of hand use. This hypothesis would be consistent with recent fMRI studies of amputees, which showed that phantom limb finger movements also match the characteristic organization found in able-bodied individuals^23,24^. That characteristic representation was identified in the sensorimotor cortex using fMRI, so 2) the BCI finger representation in PC-IP might instead match the representation of able-bodied individuals in the same brain area (anterior superior parietal lobule - SPLa; Figure 2c and Supplementary Figure 7), despite poor finger individuation outside M1/S1 at the fMRI scale^25^. Another possibility is that 3) pre-injury motor representations could have de-specialized after paralysis, such that finger activity patterns are unstructured (Figure 2c). However, this hypothesis would be inconsistent with fMRI studies of amputees’ sensorimotor cortex^23,24^. Lastly, 4) the finger movement representational structure might optimize for the statistics of the task^15,26^. Our BCI task, as well as previous experiments with participant X, involved no correlation between individual fingers. By construction, the optimal structure, requiring the least change in finger activity patterns, would represent each finger independently. In other words, the task-statistics hypothesis (4) would predict that, with BCI usage, the representational structure would converge towards the pairwise-independent representational structure (Figure 2d).

Does the finger representational structure in a tetraplegic individual match that of able-bodied individuals? We quantified the finger representational structure by measuring the cross-validated Mahalanobis distance (Methods) between each finger pair, using the firing rates from the same time window used for BCI control. The resulting RDMs are shown in Figure 2e (average across sessions) and Supplementary Figure 8 (all sessions). For visual intuition, we also projected the representational structure to two dimensions in Figure 2f, which shows that the thumb is distinct while the middle, ring, and pinky are close in neural space. We then compared the measured RDMs against the able-bodied and pairwise-independent models using the whitened unbiased RDM cosine similarity (WUC)^27^. The measured representational structure matched the able-bodied M1 representational structure significantly over the unstructured model (P = 1.8 × 10^-10^, two-tailed t-test) (Figure 2g), ruling out the de-specialization hypothesis (3). Furthermore, the fit to the able-bodied M1 structure was close to the theoretical maximum (P = 0.096, Methods), supporting our predicted hypothesis (1). Our findings were robust to different distance and model-fit metrics (Supplementary Figure 10).

The measured representational structure also matched the able-bodied M1 representational structure significantly better than the able-bodied SPLa representational structure (P = 4.5 × 10^-7^, two-tailed t-test). This match was consistent across individual able-bodied participants’ (N = 20) fMRI results, with the BCI finger representational structure matching every individual’s M1 better than their SPLa (Supplementary Figure 9). These results rule out hypothesis 2.

### Representational structure did not trend towards task optimum

Was the BCI finger representational structure consistently similar to M1? The task-optimal structure hypothesis (4) predicted that the BCI RDMs would trend to optimize for the task statistics (unstructured model, Figure 2d) as the participant performed the BCI task. However, the model fit did not trend from the M1 model towards the unstructured model (linear-model session × model interaction: *t*(3) = −0.71, one-tailed t-test P = 0.74, Bayes factor (BF) = 0.33) (Figure 3a). We also did not find evidence that the model fit started similar to SPLa (the same brain region) and trended towards M1 (linear-model session × model interaction: *t*(3) = 0.53, one-tailed t-test P = 0.32, Bayes factor (BF) = 0.73). Indeed, the representational structure was largely consistent across different recording sessions (average pairwise correlation, excluding the diagonal: r = 0.90 +/- s.d. 0.04, min 0.83. max 0.99).

**Figure 3.**
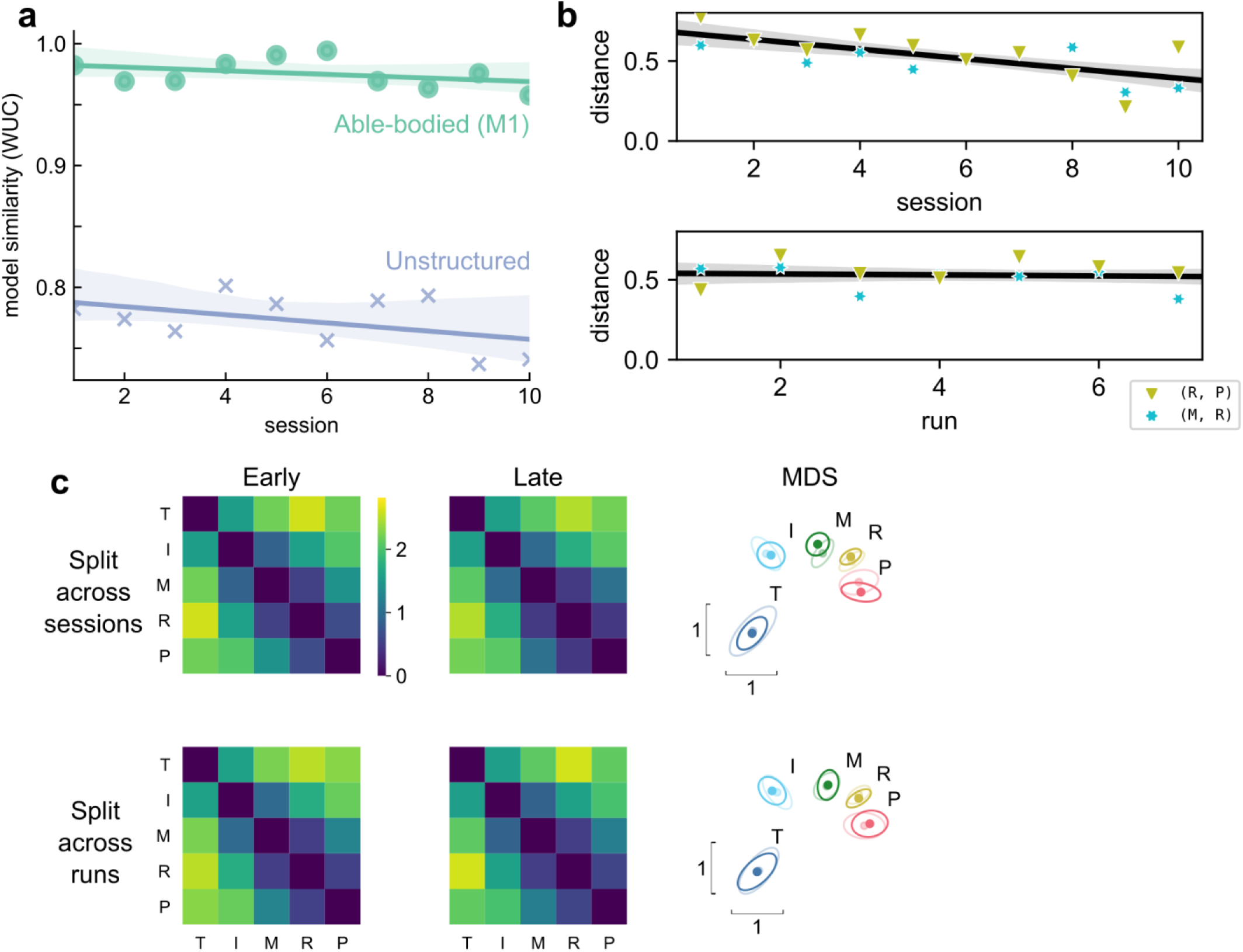
Hand representation changed minimally after weeks of BCI control. (**a**) Slope comparison shows that the model fit did not trend towards the constant model over sessions. (**b**) The distance between high-error finger pairs (middle-ring and ring-pinky) did not increase across sessions or runs (within sessions), as shown by partial regression plots. Distance metric: cross-validated Mahalanobis, averaged across runs (for the session plot) or averaged across sessions (for the run plot). The black line indicates linear regression. The gray shaded region indicates a 95% confidence interval. Each run consisted of 8 presses per finger. (**c**) Minimal change in representational structure between early and late sessions or between early and late runs. Mean RDM, when grouped by sessions (top row) or individual runs (bottom row). Grouped into early half (left column) or late half (center column). MDS visualization (right column) of early (opaque) and late (translucent) representational structures after Generalized Procrustes alignment (without scaling, to allow distance comparisons).

We considered whether learning, across sessions or within sessions, could have caused smaller-scale changes in the representational structure. The observed representational structure, where middle-ring and ring-pinky pairs had relatively small distances, was detrimental to classification performance. The majority (70%) of the online classification errors were middle-ring or ring-pinky confusions (Figure 1b). Due to these systematic errors, one might reasonably predict that plasticity mechanisms would improve control by increasing the inter-finger distances between the confused finger pairs. Contrary to this prediction, the middle-ring and ring-pinky distances did not increase over the course of the experiment (across sessions: *t*(2) = −4.5, one-tailed t-test P = 0.98, BF = 0.03; within sessions: t(2) = −0.45, one-tailed t-test P = 0.65, BF = 0.12) (Figure 3b). When analyzing all finger pairs together, the inter-finger distances also did not increase (across sessions: *t*(10) = −4.0, one-tailed t-test P = 0.999, BF = 0.01; within sessions: t(10) = −2.4, one-tailed t-test P = 0.98, BF = 0.02), as visualized by the similarity between the average early-half RDM and the average late-half RDM (Figure 3c). These analyses demonstrate that the representational structure did not trend towards the task optimum (Figure 2d), ruling out hypothesis 4.

### Finger representational structure is motor-like and then receptive-field-like

It might seem surprising that our intracortical recordings in PC-IP match the fMRI structure in M1 rather than SPLa. One possibility is that the PC-IP structure is being driven by an efference copy from M1. This hypothesis would predict temporal heterogeneity in representational structure, with an early motor-command-like component during movement initiation. To investigate this temporal evolution, we modeled the representational structure of digit movements at each time point as a non-negative linear combination^29^ of potentially predictive models (Figure 4a).

**Figure 4.**
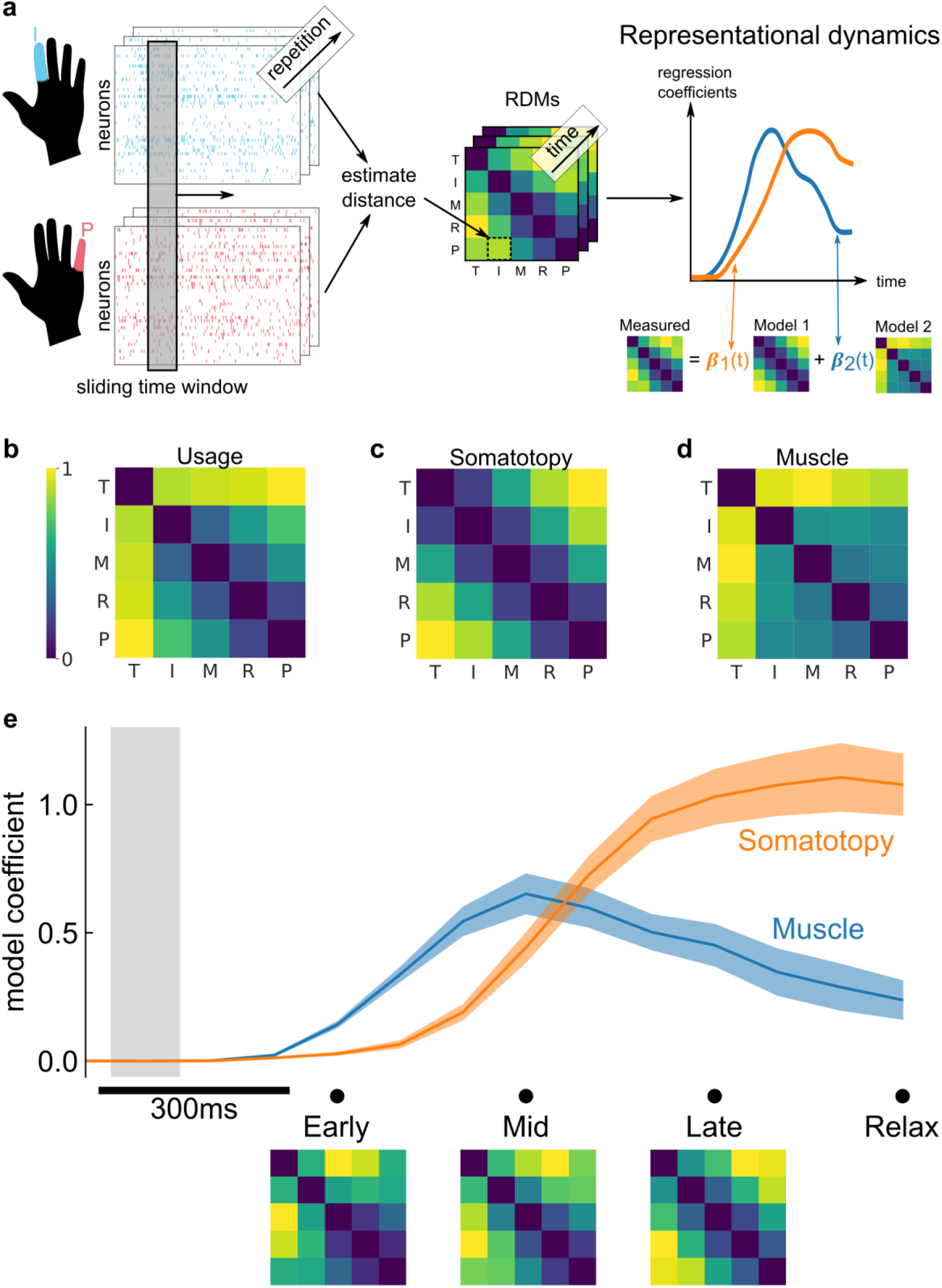
Representational dynamics analysis (RDA) dissociates neural processes over time. (**a**) RDA performs representational similarity analysis (RSA) in a sliding window across time. Here, we model the measured representational structure as a nonnegative linear combination of component model RDMs. (**b-d**) Hypothesized explanatory component RDMs: usage, muscle, and somatotopy^14^. Max-scaled to [0, 1]. (**e**) RDA of the measured RDM over time shows an early fit to the muscle model and a late fit to the somatotopy model. Confidence intervals indicate +/- s.e.m. bootstrapped across 10 sessions. Gray shaded region indicates the approximate onset time of the saccade to cue (interquartile range across trials). Difference in model start-time (170ms, Methods) was significant (P = 0.002, two-sided Wilcoxon signed-rank test). RDM snapshots (bottom, each max-scaled to [0, 1]) intuitively visualize the change in representational structure over time from muscle-like to somatotopy-like.

We considered three models^14^ that could account for representational structure: hand-usage model, muscle activation, and somatotopy. The hand-usage model (Figure 4b) predicts that the neural representational structure should follow the correlation pattern of finger kinematics during natural hand use. The muscle activation model (Figure 4c) predicts that the representational structure should follow the coactivation patterns of muscle activity during individual finger movements. The somatotopy model (Figure 4d) predicts that the representational structure should follow the spatial layout of the fingers, with neighboring digits represented similarly to each other^14,30^. At the neural population level, the somatotopy model is analogous to Gaussian receptive fields^30^.

Because the three proposed models are nearly multicollinear (max variance inflation factor = 79), we first needed to reduce the number of component models. Through a model selection procedure (Methods), we found that the hand-usage+somatotopy and muscle+somatotopy model combinations matched the data best (Supplementary Figure 12). In the main text, we present our temporal analysis using the muscle+somatotopy component models.

Figure 4e shows the decomposition of the representational structure into the muscle and somatotopy component models. The results show a dynamic structure, with the muscle model emerging 170ms earlier than the somatotopy model (P = 0.002, two-sided Wilcoxon signed-rank test). This timing difference was consistent across individual sessions (Supplementary Figure 13) and task contexts such as the calibration task (Supplementary Figure 14). Indeed, the transition from the muscle model (Figure 4c) to the somatotopy model (Figure 4d) is visually apparent when comparing the average RDMs at 600ms (muscle-model-like) and 1200ms (somatotopy-model-like) (Figure 4e).

These temporal dynamics were robust to our feature selection procedure, demonstrating a similar timing difference for the hand-usage+somatotopy combination (Supplementary Figure 14), or even all three models (with somatotopy lagging muscle and hand-usage) when regularizing model fits. This approach does implicitly group the muscle and hand-usage component models as motor production models (Discussion), as they are highly correlated (r = 0.90).

## Discussion

### Neural prosthetic control of individual fingers using recordings from PC-IP

We found that participant X could robustly control a single-finger neural prosthetic in a variety of contexts (Figure 1, Supplementary Figure 3, Supplementary Figure 4), despite years of paralysis. Our BCI classification accuracy exceeded the previous best online finger control in humans^31^. Furthermore, although previous studies had shown that the anterior intraparietal area (AIP) of PPC is involved in whole-hand grasping^19,32,33^, our work is the first to demonstrate individual finger representation in PPC (Supplementary Figure 5).

### Comparing single-neuron and fMRI recordings

fMRI studies have shown that the hand area of M1/S1 exhibits a characteristic functional organization, predicted by the natural statistics of use^14^. We hypothesized that fMRI recordings of M1 could provide a reasonable model for two reasons. First, we found that PC-IP neurons encode individual finger movements (Supplementary Figure 5), but the sensorimotor cortex is the only cortical area to show consistent finger representation at the voxel scale^25^. Second, PPC is bidirectionally connected to the motor cortex and receives an efference-copy of M1 output. Our single-neuron recordings closely matched the M1 fMRI representational structure, reaching similarities close to the noise ceiling (Figure 2g).

We also compared our PC-IP neuronal recordings with fMRI recordings of SPLa, the region of interest (ROI) that anatomically best corresponds with the location of the array implant. While the PC-IP recordings fit the RDM of SPLa better than the task-optimal structure, they did not fit as well as the M1 organization (Figure 2g). Why wasn’t there a stronger match with fMRI recordings of SPLa? Although fMRI and neuronal recordings can produce similar RDMs^22^, the RDMs can also differ^34^ when representations are organized heterogeneously at different scales^35^. For example, the representational structure of finger flexion and extension differs between fMRI and single-neuron recordings in M1. This is thought to result from flexion/extension neural populations that cluster spatially or share inputs^34^. Likewise, we found that PC-IP finger encoding is heterogenous at the single-neuron level (Supplementary Figure 5), even while the posterior parietal cortex lacks a clear finger topography^25,36,37^. SPLa fMRI finger RDMs are thus more variable and may be difficult to use as a model (Supplementary Figure 7).

### Matching representations in able-bodied and paralyzed participants

We discovered a match in finger movement representations between able-bodied and paralyzed participants. This match suggests that pre-injury finger representations have been preserved even after years of paralysis (C3-C4 AIS-A spinal cord injury). To control neural prosthetic fingers, PX could reactivate these latent motor representations.

Our finding of preserved representations adds to an evolving and multifaceted understanding of plasticity after sensorimotor loss. Early studies by Merzenich and colleagues showed that S1 reorganized after amputation, with intact body parts invading the deprived cortex^6–8^. However, the authors also recognized that the amputated body part might persist in latent somatosensory maps. Preserved, latent somatosensory representations were later proven correct by recent studies of amputation^9,23,24,38^ and even paralysis^39,40^. A recent motor cortex BCI study^41^ also found a similar pattern of finger classification errors to the characteristic able-bodied structure (Figure 1b and Figure 2b), hinting that the motor cortex of tetraplegic participants may also match that of able-bodied participants. Fewer studies have investigated sensorimotor plasticity beyond M1/S1, but our results in PC-IP indicate that higher-order regions can remain surprisingly preserved after paralysis.

The significance of cortical reorganization has long been discussed in the field of BCIs, particularly when deciding where to implant electrodes. If, as previously thought, sensory deprivation drives cortical reorganization and any group of neurons can learn to control a prosthetic^42,43^, the specific implant location would not affect BCI performance. Our results and others^2,9,12,23,24,38^ suggest that such a stance is oversimplified. Although experience does shape neural organization^6,14,24^, representations may be remarkably persistent once formed^24^. Thus, to enable intuitive control for prosthetic users, neural prosthetics will benefit from tapping into the preserved, natural^12^ movement repertoire of motor areas.

### Consistent representational structure across sessions

We found that the BCI representational structure changed minimally over weeks (Figure 3) despite the structure’s contributions to misclassification (Figure 1b). While PX was motivated to perform well and was anecdotally understood which finger pairs the online classifier was confusing, she could not increase the neural distance between fingers. Our observed lack of learning matches results from single-session learning studies, where neural learning is limited to the intrinsic neural manifold^12,44–46^.

A study of long-term learning using artificial perturbations found that rhesus monkeys can learn to generate novel neural patterns after about 8 sessions^47^. The difference with our results could indicate that PX needed a stronger learning pressure, or perhaps the monkeys learned to use an alternative cognitive strategy like moving multiple effectors. Here, the participant did not attempt any alternative movements. Additionally, because we were interested in understanding the natural finger representation, we did not artificially perturb the BCI mapping to increase learning pressure. The error structure was instead a product of the natural representational structure. We expect future studies will further clarify how and when BCI learning can occur, as ours is only the first to investigate long-term population learning in a human BCI framework.

### Representational dynamics consistent with PPC as a forward model

Based on previous studies of the posterior parietal cortex, different neural processes could be salient at different times during movement. PPC is thought to maintain an internal model of the body^48–51^. As such, PPC receives efference copies of motor command signals and delayed multimodal sensory feedback. The internal model role predicts that PPC houses multiple functional representations, each engaged at different time points of motor production.

We performed a time-resolved version of representational similarity analysis to dissociate neural processes over short time scales (hundreds of milliseconds, Figure 4). Our temporal analysis showed a consistent ordering: early emergence of the muscle or hand-usage components followed by the somatotopy model.

We found consistent temporal results when using either the muscle or hand-usage component models (Figure 4 and Supplementary Figure 14), as hand-usage and muscle activation patterns are strongly correlated for individual finger movements^52^. Therefore, here we group these models under the single concept of motor production. In the future, more complex multi-finger movements^14^ would help distinguish between muscle and hand-usage models.

The somatotopy model derived from a simple-topography model predicts that neighboring digits will have similar cortical activity patterns^14^. However, at the neural population level, the same representational structure is more analogous to Gaussian receptive fields^30^. Gaussian receptive fields have been useful tools for understanding digit topographies within the sensorimotor cortex^30,53^. In another study with participant X, we found that the same recorded population encodes actual touch^54^ with a Gaussian-like receptive fields. Based on these results, here the somatotopy model can be thought of as a sensory-consequence model. However, because PX has no sensation below her shoulders, we interpret the somatotopy model as the preserved prediction of sensory consequences of a finger movement. These sensory outcome signals could be the consequence of internal computations within the PPC or could come from other structures important for body-state estimation, such as the cerebellum^51^.

The 170ms timing difference we found roughly matches the 60ms + 60ms delay between feedforward muscle activation and somatosensory afferents^55,56^. In able-bodied individuals, PPC is thought to maintain a state estimate for motor production, which would integrate motor planning, production, and predicted-sensory-outcome signals at such a timing^48–51^. The matching timing, even during BCI control, provides further evidence that the recorded motor circuits have preserved their functional role.

### Preserved motor representations after paralysis

A persistent question in neuroscience has been how experience shapes the brain, and to what extent existing neural circuits can be modified. In the first human BCI study of neural-population plasticity, we found that the brain’s motor circuits seem remarkably stable even after severe injury. An interesting question for future studies is whether more complex motor skills, such as handwriting^57^, remain preserved through injury. These findings will continue to influence the design of neural prosthetics and help to restore motor abilities to people with paralysis.

## Materials and methods

### Data collection

#### Study participant

The study participant X (abbreviation: PX) has a AIS-A spinal cord injury at cervical level C3-C4 that she sustained approximately ten years before this study. PX cannot move or feel her hands. As part of a BCI clinical study (ClinicalTrials.gov identifier: NCT01958086), PX was implanted with two 96-channel Neuroport Utah electrode arrays (Blackrock Microsystems model numbers 4382 and 4383). She consented to the surgical procedure as well as to the subsequent clinical studies after understanding their nature, objectives, and potential risks. All procedures were approved by the California Institute of Technology, Casa Colina Hospital and Centers for Healthcare, and the University of California, Los Angeles Institutional Review Boards.

#### Implant methodology and physiological recordings

The electrode array used here was implanted over the hand/limb region of the left PPC at the junction of the intraparietal sulcus (IPS) with the postcentral sulcus (PCS) (Supplementary Figure 1). We previously^19,45,58^ referred to this brain area as the anterior intraparietal area (AIP), a region functionally defined in non-human primates (NHPs). In this report, we use anatomical characteristics to name this brain area, denoting it the postcentral-intraparietal area (PC-IP). More details regarding the methodology for functional localization and implantation can be found in ^45^.

#### Neural data preprocessing

Unit activity was detected by thresholding the waveform at −3.5 times the root-mean-square, after high-pass filtering (250Hz cut-off) the full-bandwidth signal. Single and multiunit activity was sorted using k-medoids clustering using the gap criteria^59^ to determine the total number of neural clusters. Clustering was performed on the first *n* ∈ {2, 3, 4} principal components, where *n* was selected to account for 95% of waveform variance.

### Experimental setup

#### Recording sessions

Experiments were conducted in 2-3 hour recording sessions at Casa Colina Hospital and Centers for Healthcare. All tasks were performed with PX seated in her motorized wheelchair with her hands resting prone on the armrests. PX viewed cues on a 27-inch LCD monitor that occupied approximately 40 degrees of visual angle. Stimulus presentation was controlled by the psychophysics toolbox^60^ for MATLAB (Mathworks).

The data were collected on 9 days over 6 weeks. Almost all experiment days were treated as individual sessions (i.e., the day’s recordings were spike-sorted together). The second experiment day was an exception, with data being recorded in a morning period and an afternoon period with a sizable rest period in between. To reduce the effects of recording drift, we treated the two periods as separate sessions (i.e., spike-sorted each separately), for a total of 10 sessions. Each session can thus be considered a different resampling of a larger underlying neural population, with some unique neurons and some repeated neurons. We did not re-run the calibration task for the afternoon session, resulting in 9 sessions of the calibration task for Supplementary Figure 3b.

#### Calibration task

At the beginning of each recording session, a reaction-time finger flexion task (denoted “calibration task” in the Results) was performed to train a finger classifier for online control during subsequent runs of the primary task. On each trial, a letter appeared on the screen (e.g., “T” for thumb). The participant was instructed to immediately flex the corresponding finger on the right hand (contralateral to the implant), as though striking a key on a keyboard. Conditions were interleaved in a pseudorandom order such that each condition was performed once before repetition.

The classifier was then calibrated according to the Finger Classification section. The calibration task did not have a No-Go condition, so the firing rates during the intertrial interval (ITI) were used instead to train the classifier’s No-Go prediction.

#### Finger flexion grid task

In the primary task, movement cues were arranged in a 3 x 4 grid of letters on the screen. Each screen consisted of two repetitions each of T (thumb), (index), M (middle), R (ring), P (pinky), and X (No-Go) arranged randomly on the grid. Every three seconds (i.e., each trial), a new cue was randomly selected with a crosshairs indicator, which was jittered randomly to prevent letter occlusion. Each cue was selected once (for a total of 12 trials) before the screen was updated to a new arrangement. Each block consisted of 3-4 screens.

On each trial, the participant was instructed to immediately saccade to the cued target and fixate, then attempt to flex the corresponding finger. During both movement and No-Go trials, the participant was instructed to fixate on the target at least until the visual classification feedback was shown. The randomized cue location was intended to investigate whether cue location affects movement representations.

The classifier decoded the finger movement on each trial and presented its prediction via text feedback 1.5 seconds after the cue presentation.

#### No-movement control task

The control task was similar to the primary task, except the subject was instructed to saccade to each cued letter and fixate without attempting any finger movements. No classification feedback was shown.

### Statistical analysis

#### Unit selection

Single-unit neurons were identified using the k-medoids clustering method, as described in the Neural Data Preprocessing section. Analyses in the main text used all identified units, regardless of sort quality. With spike-sorting, there is always the possibility that a single waveform cluster corresponds to activity from multiple neurons. To confirm that potential multi-unit clustering did not bias our results, we repeated our analyses using only well-isolated units (Supplementary Figure 15).

Well-isolated single units were identified using the L-ratio metric^61^. The neurons corresponding to the lowest third of L-ratio values (across days) were selected as “well-isolated.” This corresponded to a threshold of 10^-1.1^ dividing well-isolated single units and potential multi-units (Supplementary Figure 15).

#### Single-unit tuning to finger flexion

To calculate significance for each neuron (Supplementary Figure 5), we used a two-tailed t-test comparing each movement’s firing rate to the No-Go firing rate. A neuron was considered significantly tuned to a movement if P < 0.05 (after FDR correction). We also computed the mean firing rate changes for each condition. If a neuron was significantly tuned to at least one finger, we denoted the significant finger with the highest mean firing rate change as the neuron’s “best finger.” Discriminability index (d’, RMS standard deviation) was computed between the No-Go mean firing rate, as the baseline, and the mean firing rate during each condition.

Units were pooled across all 10 sessions. Units with mean firing rates less than 0.1 Hz were excluded to minimize sensitivity to discrete spike-counting.

#### Finger classification

Finger classification was performed using linear discriminant analysis (LDA) with diagonal covariance matrices^62^; diagonal LDA is also equivalent to Gaussian Naive Bayes with a shared covariance matrix.

To calibrate the in-session classifier, LDA was fit to the binned threshold crossings in a 1-second time window of trials of the calibration task, where the window lag was chosen to maximize the cross-validated classification accuracy for that trial block. Electrodes with mean firing rates less than 1 Hz were excluded to prevent low-firing rate discretization effects. This classifier was then used in subsequent online control in the main task.

During online control of the finger flexion grid task, classification features were constructed using the binned threshold crossings from each electrode in the time window [0.5, 1.5] seconds after cue presentation. The window start-time was chosen based on the estimated saccade latency in the first experimental session. The saccade latency was estimated by taking the median of the time the subject took to look > 80% of the distance between targets. The analysis window was a priori determined to be 1 second; this choice was further supported by a sliding window analysis confirming accurate finger decoding up to 1.6 seconds.

Offline classification accuracy was computed using leave-one-out cross-validation. We used features from the same time window as the online control task. However, offline analyses used spike-sorted firing rates instead of electrode threshold crossings.

To visualize aggregate confusion matrices, confusion matrix counts were summed across recording sessions, then normalized by row (true label) to display ratios. This is equivalent to pooling trials across sessions to create a single confusion matrix. Reported classification accuracies aggregate trials over all sessions. Reported standard deviations are over sessions, weighted by the number of trials in each session.

#### Distance measure

A robust variant^63^ of the cross-validated (squared) Mahalanobis distance was used to measure the dissimilarity between neural patterns for each pair of fingers (*j, k*). The cross-validated Mahalanobis distance is calculated across independent partitions A and *B*, each with respective noise covariance Σ and measured activity patterns (*b_j_, b_k_*) :

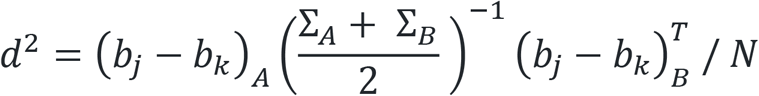

*N* normalizes for the number of neurons, so the units of *d*^2^ are (*unitless*^2^/*neuron*).. Here, we split the trials into 5 independent partitions and averaged the distance estimate across all permutations of partition pairs (*A, B*). The cross-validated Mahalanobis distance measures the separability of multivariate patterns and is the continuous analog to LDA classification accuracy. Cross-validation ensures the (squared) distance estimate is unbiased; 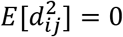 when the underlying distributions are identical^64^.

The noise covariance matrix, Σ, was used to normalize noise across different neurons. Σ was estimated from the data independently for each cross-validation fold. We regularized Σ towards a diagonal covariance matrix^65^ to guarantee that the estimate was invertible.

Because RDMs are symmetric and the dissimilarity metric is unbiased, only the RDM’s lower triangular values were used for subsequent model-fitting procedures.

For comprehensiveness, we also compute RDMs using a cross-validated version of the Poisson symmetrized KL-divergence^63^ in Supplementary Figure 10.

#### Representational models

The M1 model for Figure 2g was taken from the right-hand 7T scans of Ejaz et al.^14^. The SPLa model was taken from the pre-experiment, right-hand 3T scans of Kieliba et al.^28^. To fairly compare the reliability of finger RDMs in M1 and SPLa, Supplementary Figure 7 uses 3T fMRI data from the same subjects and scans^28^. Similar results for Figure 2g were found using M1 data from both studies^14,28^.

The hand usage, muscle, and somatotopy models were taken from ^14^. The natural movement statistic model was constructed using the velocity time series of each finger’s MCP joint during everyday tasks. The muscle activity model was constructed using EMG activity during single- and multi-finger tasks. The somatotopic model is based on a cortical sheet analogy and assumes that finger activation patterns are linearly spaced Gaussian kernels across the cortical sheet. Further modeling details are available in the methods section of ^14^.

#### Comparing representational structures

The rsatoolbox Python library^63^ was used to perform representational similarity analysis^20^ between the models and our data.

To quantify model fit, we used the whitened unbiased RDM cosine similarity (WUC) metric, as is recommended for quantitative prediction models^27^. Unlike correlation metrics, which are 0 when the model RDM and data RDM have no systematic relationship, WUC > 0 as long as the model predicts positive distances between neurally distinct conditions. Therefore, WUC values are often larger than the corresponding correlation values or are even close to 1. Instead of comparing against 0, WUC values can be interpreted by comparing against a baseline. The baseline is usually chosen to be a null model where all conditions are equidistant pairwise (and would thus correspond to a 0-correlation). Here, our BCI task structure is equivalent to the null model, because the fingers were cued in random order and individually in a counterbalanced manner. The task thus had zero correlation between conditions. For comprehensiveness, we also show model fits using whitened Pearson correlation in Supplementary Figure 10. Whitened Pearson correlation is a common alternative to WUC^27^.

To estimate the noise ceiling, we calculated the average similarity of each individual-session RDM with the mean RDM across sessions^66^. This value is a slight overestimate of the true noise ceiling. To calculate a lower-bound estimate of the noise ceiling, we calculated the average similarity of each individual-session RDM with the mean RDM across all other sessions (i.e., excluding that session). The area between the lower and upper bounds was shaded as the noise ceiling region.

#### Measuring changes in the representational structure

To assess the effect of BCI task experience on the inter-finger distances, we performed a linear model analysis with predictors of session index, within-session run-block index, and finger pair. Cohen’s *f*^2^ for each predictor was calculated using 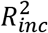, the increase in *R*^2^ from the finger-pair-only model to the model including both the specified predictor and the finger pair.

Bayesian tests were computed using the R package BayesFactor^67^ with default priors. To test each directional hypothesis, we sampled (N = 10^6^) from the posterior of the corresponding non-directional model, calculated the proportion of samples that satisfied the directional hypothesis, and divided by the prior odds^68^.

#### Linear combinations of models

We also compared the data RDMs with nonnegative linear combinations of multiple model RDMs. To prevent overfitting, we cross-validated the model fits by fitting the mixture coefficients on all-but-one sessions and evaluated the model fit on the left-out session. To estimate model-fit uncertainty, we bootstrapped RDMs (sessions) over the cross-validation procedure^63^. We then selected the best model using the “one-standard error” rule^69^, choosing the simplest model within one standard error of the best model fit.

#### Representational dynamics analysis

To investigate how the finger movement representational structure unfolds over time, we followed Kietzmann et al.^29^ in using a time-resolved version of representational similarity analysis (Figure 4a). At each point in time, we computed the instantaneous firing rates by binning the spikes in a ±100ms time window centered at that point. These firing rates were used to calculate cross-validated Mahalanobis distances between each pair of fingers and generate an RDM at each time point.

The temporal sequence of RDMs constitutes an RDM “movie” (size [*n_digits_, n_digits_, n_timepoints_*]) that visualizes the representational trajectory across the trial duration. RDM movies were computed separately for each recording session. Single-time snapshots (Figure 4e) show RDMs averaged across sessions.

At each time point, we decomposed the data RDM into the component models using nonnegative least squares. Because the component models are not completely orthogonal, component models were limited to the subsets chosen in the model reduction step. Each component RDM was normalized by its vector length (ℓ_2_-norm) before decomposition to allow comparison between coefficient magnitudes. To compute confidence intervals, we bootstrapped RDMs across days, calculated the average RDM of each bootstrap sample, and then decomposed the corresponding average RDM into mixture coefficients.

We computed the start-time of each model component as the time at which the corresponding mixture coefficient exceeded 0.2 (about 25% of the median peak-coefficient across models and sessions).

### Data availability

Data will be deposited in the BRAIN Initiative DANDI Archive before publication.

### Code availability

The custom analysis code will be made available at https://github.com/AndersenLab-Caltech/fingers_rsa before publication.

## Supporting information

Supplemental Figure 4 - Controlling a virtual hand

## Author contributions

T.A., C.G., and R.A.A. designed the study. T.A and C.Y.Z. developed the experimental tasks and collected data. C.G. and T.A. analyzed the results. C.G. and T.A. interpreted results. C.G. and T.A. wrote the paper. E.R.R. provided experimental facilities and coordinated with Casa Colina Hospital and Centers for Healthcare. N.P. performed the surgery to implant the recording arrays.

## Acknowledgments

We thank participant X for her dedicated participation in the study. Kelsie Pejsa and Viktor Scherbatyuk for administrative and technical assistance. Paulina Kieliba, Elena Amoruso, and Tamar Makin for sharing their M1/SPLa fMRI data. Tamar Makin for her comments on the manuscript. Jörn Diedrichsen and Spencer Arbuckle for publicly sharing their M1 data and models.

## Supplementary figures

**Supplementary Figure 1.**
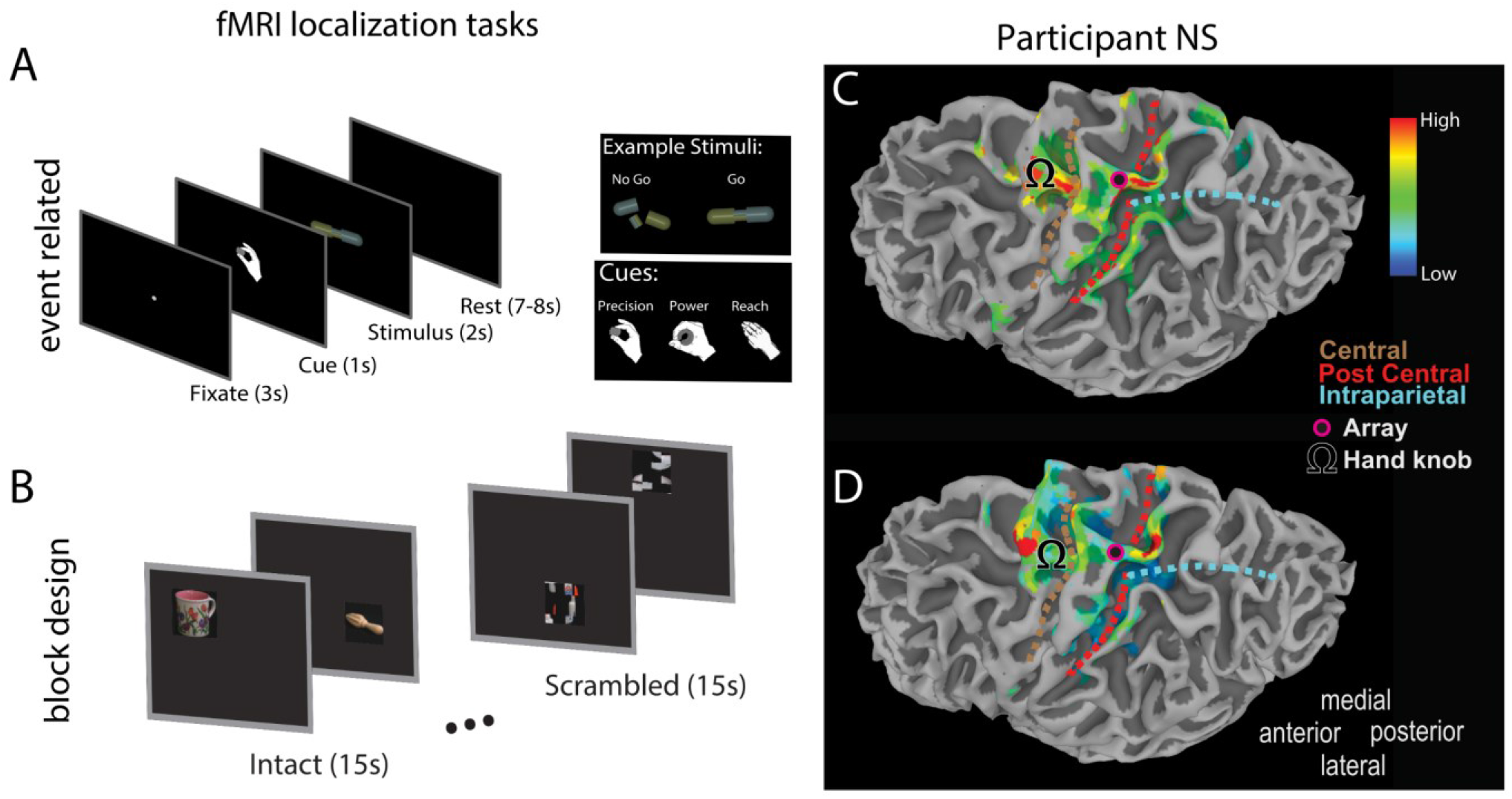
Multielectrode array implant location. Figure and text adapted from Aflalo et al.^70^ (CC BY-NC 4.0). We used fMRI to identify cortical regions involved in imagined reaching and grasping actions. We performed two complementary tasks to ensure activation was robust across paradigms. (**a**) Event-related task design. Following an intertrial interval, the subject was cued to perform a specific imagined movement (precision grasp, power grasp, or reach without hand shaping.) Following the cue, a cylindrical object was displayed. If the object was intact, the subject imagined performing the cued movement. If the object was broken, the subject withheld movement. (**b**) Block task design. Eight blocks were presented for 30 seconds per run. During the first 15 seconds of each block, common objects were presented every three seconds in varying spatial locations. Before each run, the subject was instructed to either imagine pointing at, imagine reaching and grasping, or look naturally at the object. During the last 15 seconds of each block, scrambled images were presented and the subject was instructed to guess the identity of the object. (**c**) Statistical parametric map showing voxels with significant activity for grasping (“Go” versus “No-Go”) (p < 0.01, FDR-corrected), based on task (a). Array location and cortical landmarks as depicted in the legend. (**d**) Statistical parametric map showing voxels with significant activation (P < 0.01, FDR-corrected) for grasping versus looking, based on task (b).

**Supplementary Figure 2.**
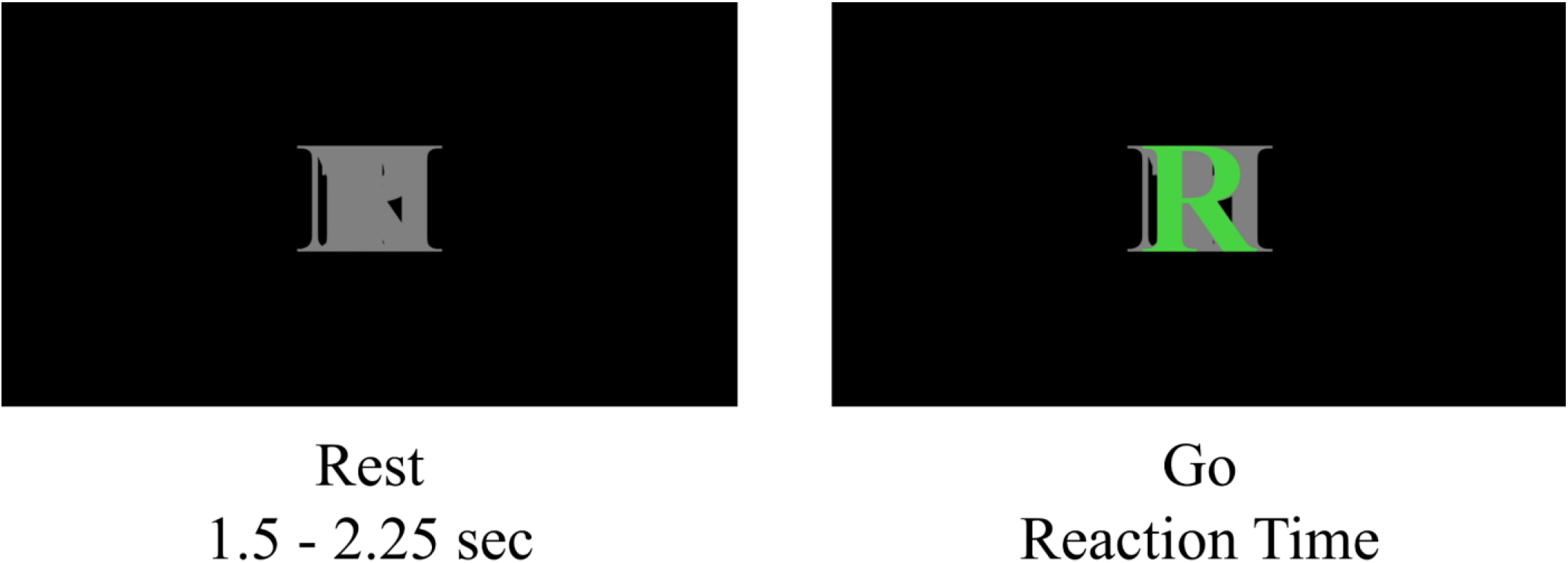
Calibration task. Task structure, single trial. Each trial consisted of an intertrial interval (ITI) and a reaction-time Go phase. During the Go phase, green text specified which digit to flex. All letters were overlaid in gray to minimize visual differences between ITI and Go phases. (Legend) T = thumb, I = index, M = middle, R = ring, P = pinky

**Supplementary Figure 3.**
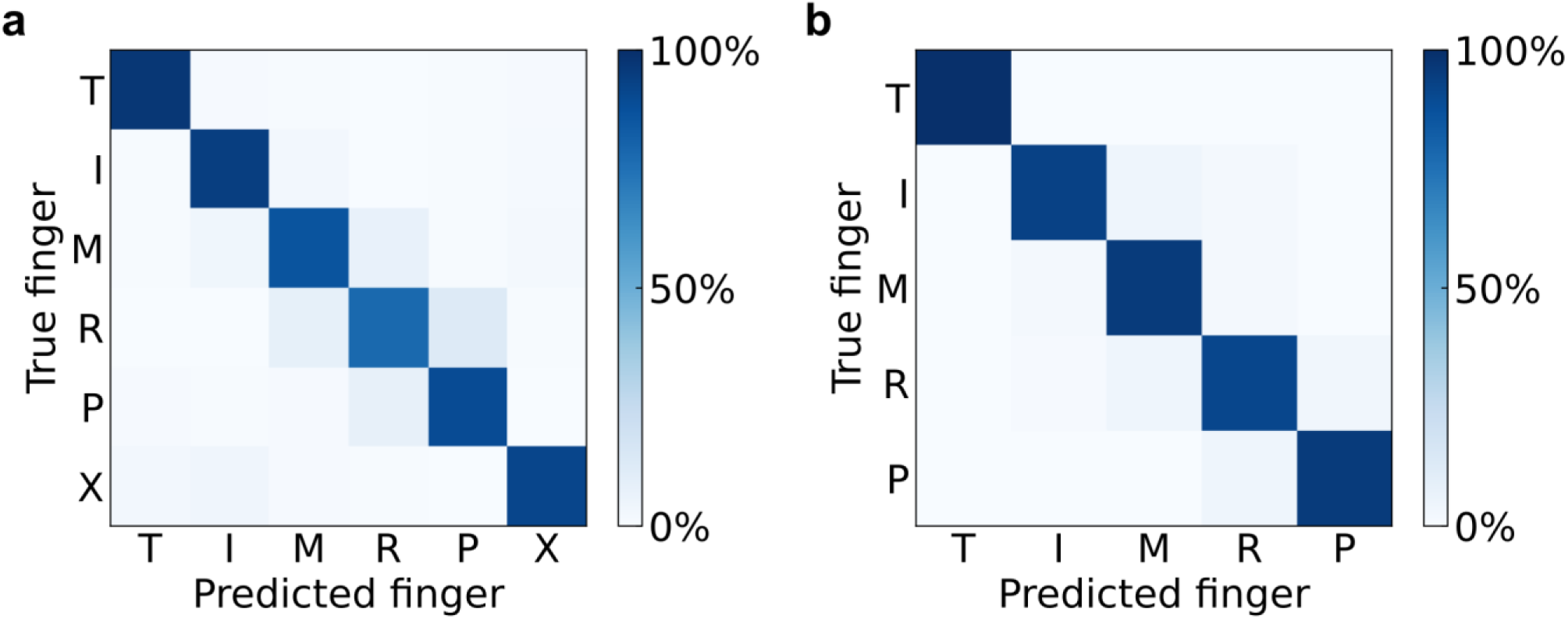
Robust cross-validated finger classification during main and calibration tasks. (**a**) Confusion matrix of offline finger classification, cross-validated within single sessions. 4080 trials of the main task aggregated over 10 sessions. (**b**) Confusion matrix of offline finger classification, cross-validated within single sessions. 530 trials of the calibration task aggregated over 9 sessions. (Legend) T = thumb, I = index, M = middle, R = ring, P = pinky, X = no movement. Each entry (*i*, *j*) in the matrix corresponds to the ratio of movement *i* trials that were classified as movement *j*.

**Supplementary Figure 4.**
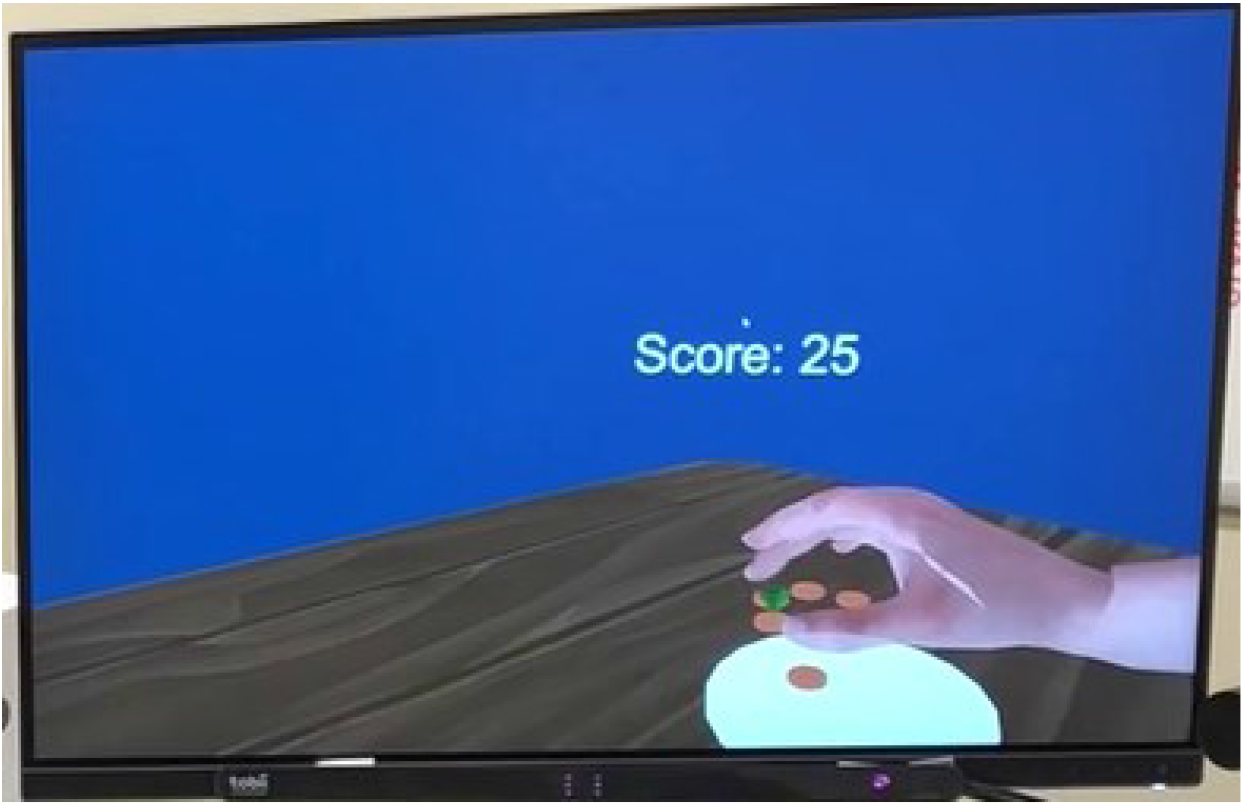
Example BCI control of a virtual reality hand. Video attached separately. Using a BCI, participant X controls the individual fingers of a virtual reality hand. She views a virtual hand, table, and cues through an Oculus headset. Similar to the main finger movement task, she acquires green jewels by pressing the corresponding finger and avoids red amethysts by resting. Green jewels disappear when the correct finger is classified (or at the start of the next trial, if incorrectly classified). The screen copies the view PX sees through the Oculus headset.

**Supplementary Figure 5.**
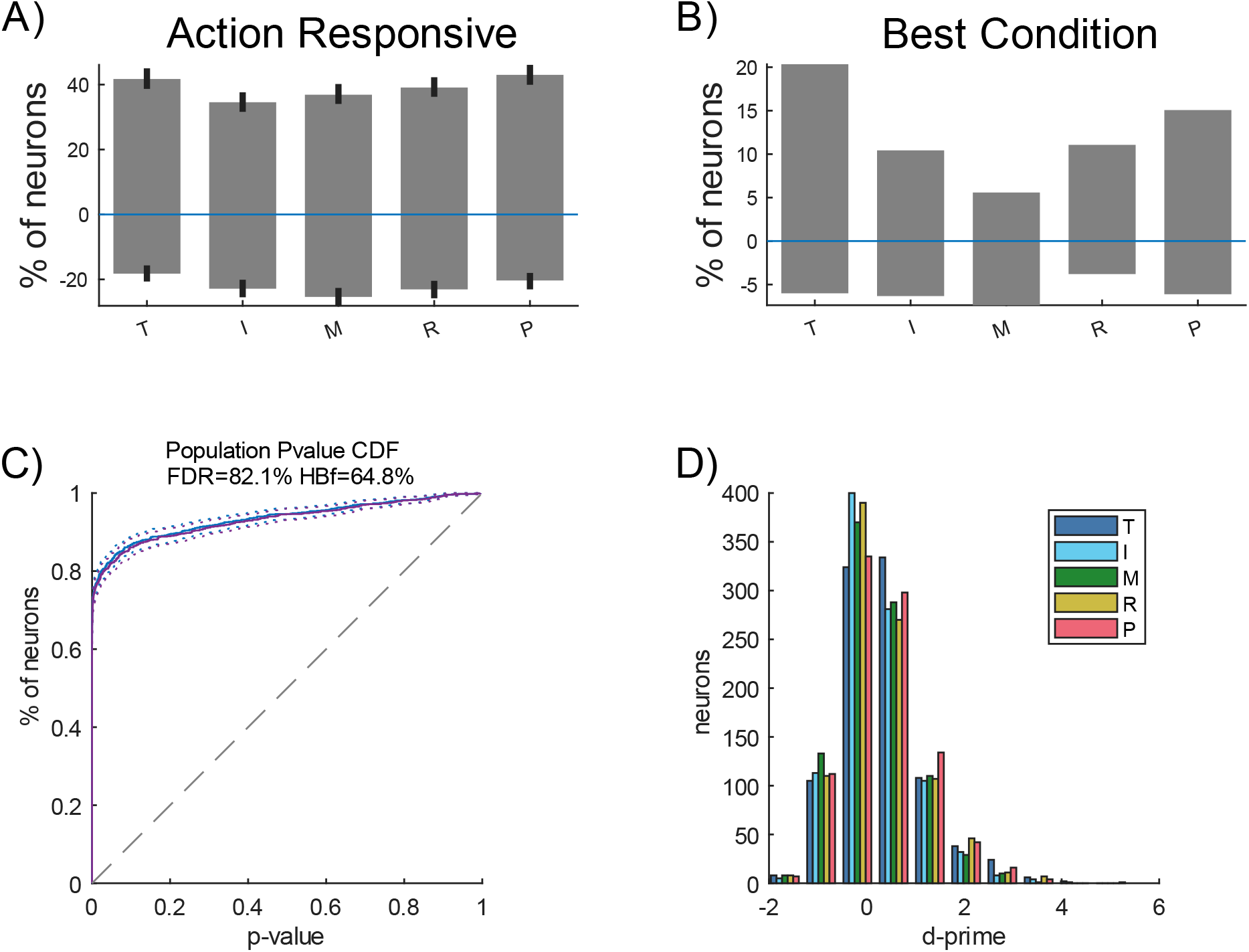
Single-neuron encoding of individual fingers. All five fingers of the right (contralateral) hand were encoded within the population during movement execution. (**a**) Percentage of the population tuned significantly (P < 0.05, FDR-corrected) to flexion of each digit. Positive percentages indicate neurons that increased firing rate during digit movement and negative percentages bar indicate neurons that decreased firing rate. Error bars indicate a 95% bootstrap confidence interval. (**b**) Percentage of the population tuned best to each digit. (**c**) Cumulative distribution function of the population’s tuning significance p-values. (**d**) Histogram of d’ (discriminability index) values across neurons. (**a-d**) Neurons were pooled across sessions.

**Supplementary Figure 6.**
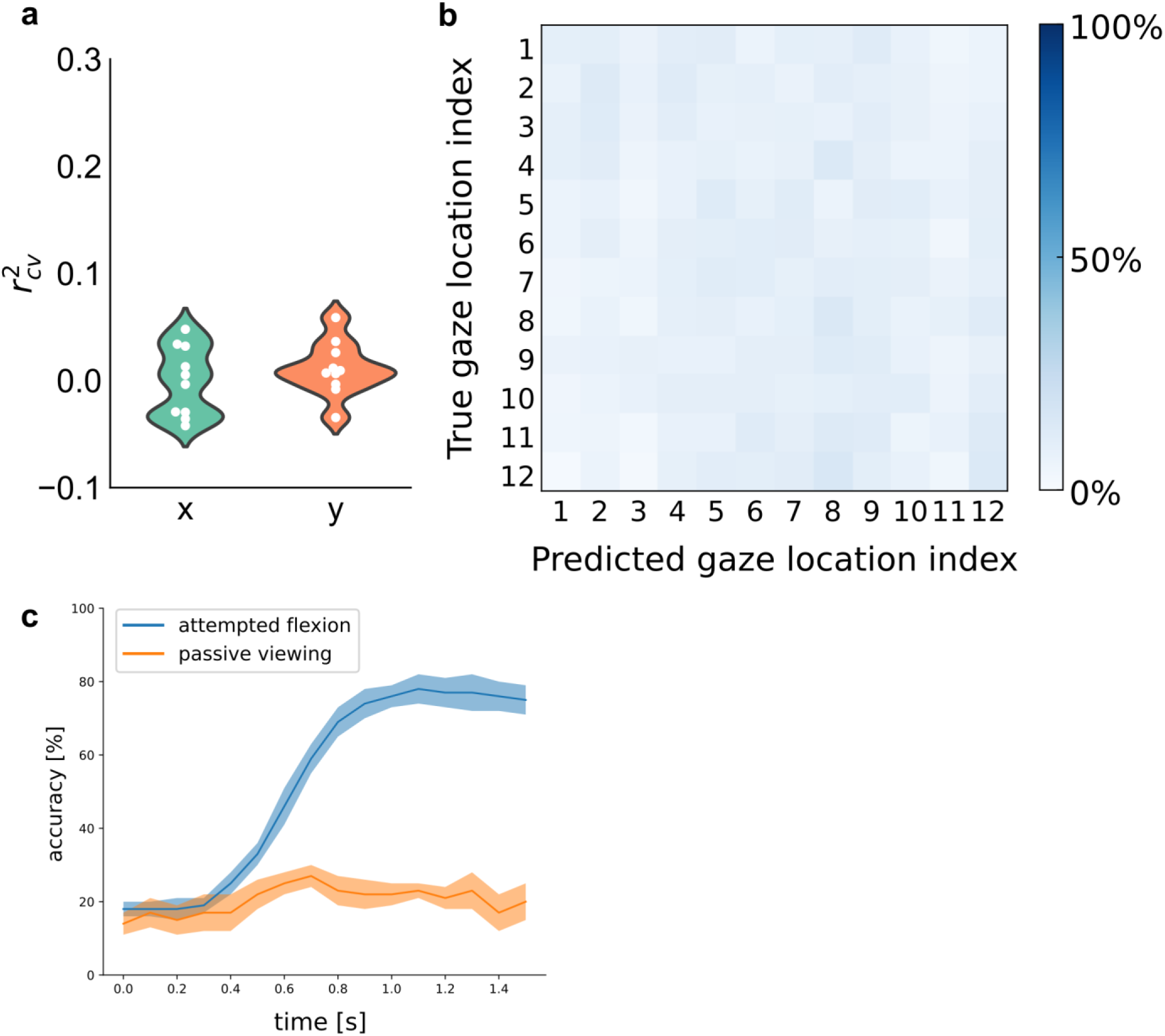
Gaze location did not affect finger decoding during the attempted-movement period. (**a**) Linear regression could not decode target location [x, y] coordinates from the neural activity during the attempted-movement period. Violin plot shows that cross-validated regression r^2^ values are close to 0 across sessions, with each circle marking a single session. (**b**) Cross-validated confusion matrix for cue location: a linear classifier (diagonal LDA) could not classify the gaze location from neural activity during the attempted-movement period. (**c**) Cross-validated classification accuracy for main and control tasks: a linear classifier (diagonal LDA) could not classify finger movements from neural activity during passive observation (orange) of the digit flexion task. Sliding bin width: 200ms. The shaded region indicates +/- s.e.m. (6 sessions passive viewing, 10 sessions attempted flexion).

**Supplementary Figure 7.**
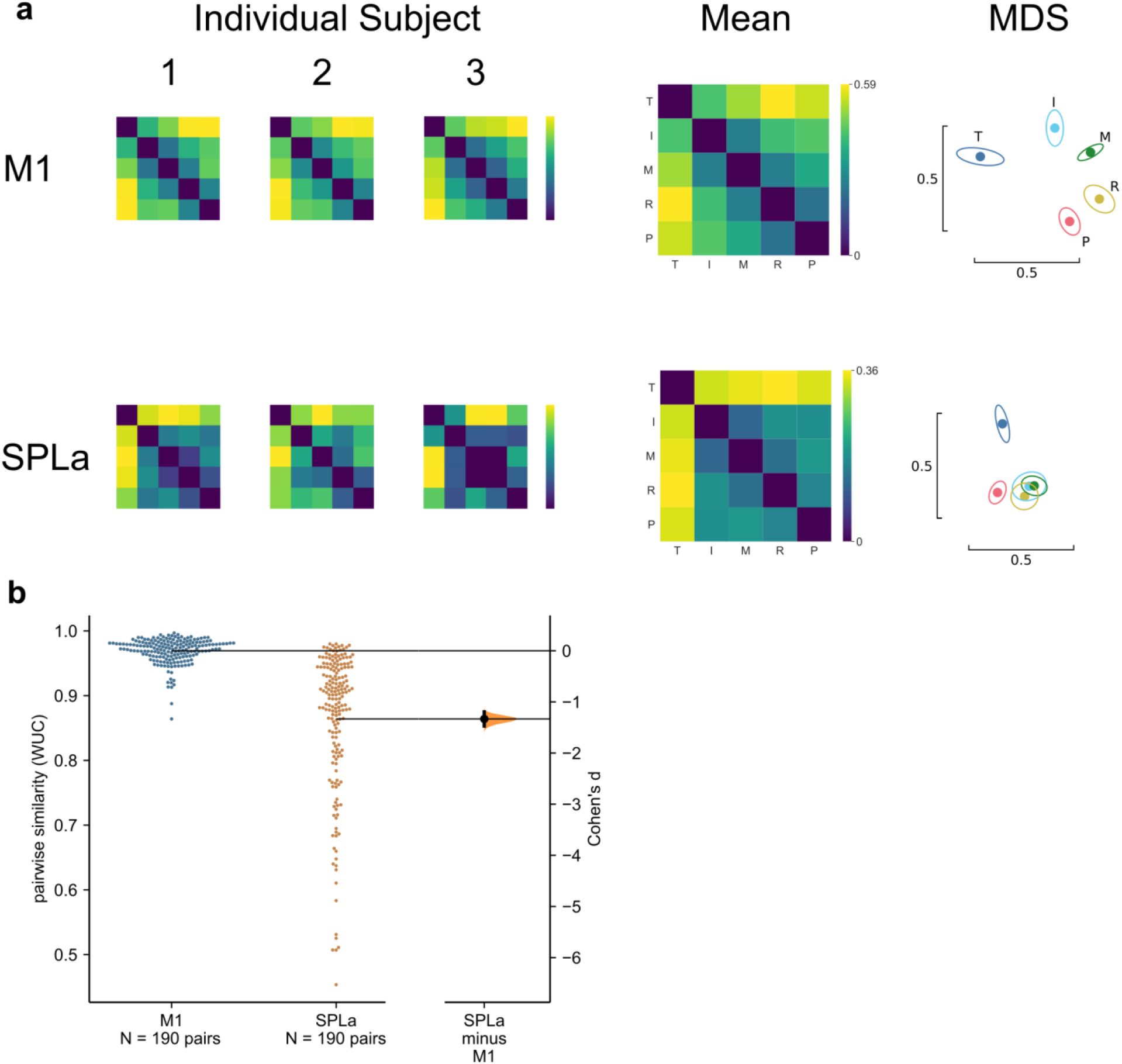
fMRI representational structure for finger movements, from Kieliba et al.^28^. (**a**) fMRI representational structure for 3 individual subjects and the group mean (N = 20). Intuitive visualization using multidimensional scaling (MDS) and Generalized Procrustes alignment (without scaling); ellipses show mean +/- s.d. across subjects. Regions of interest (ROIs): primary motor cortex (M1, top row) and anterior superior parietal lobule (SPLa, bottom row). SPLa ROI was defined in ^71^. (**b**) Finger RDMs are more consistent across subjects in M1 than in SPLa, as shown by the Gardner-Altman estimation plot^72^ of the average WUC between pairs of subject RDMs (N = 190 pairs between 20 subjects). Each circle on the swarm plot (left) marks the similarity for a pair of subjects. Horizontal black lines mark the mean of each ROI. The curve (right) indicates the resampled (N = 5000) distribution of the effect size between ROIs, as measured by Cohen’s d. Unpaired Cohen’s d between M1 and SPLa: −1.33 (95% CI: [−1.47, −1.19]).

**Supplementary Figure 8.**
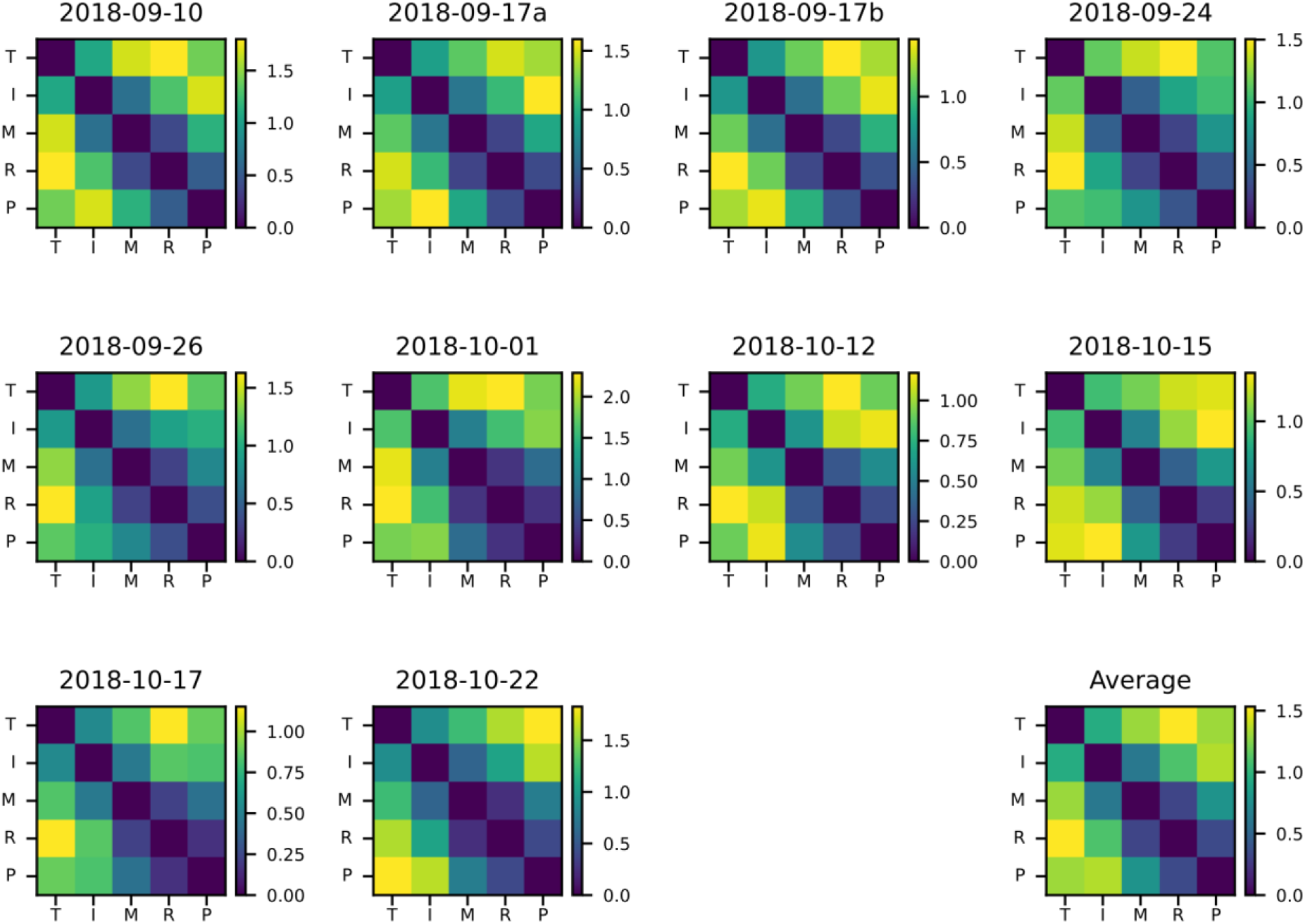
Individual representational dissimilarity matrices for each session. Representational dissimilarity matrices across all sessions, using the cross-validated Mahalanobis distance. Related to Figure 2e.

**Supplementary Figure 9.**
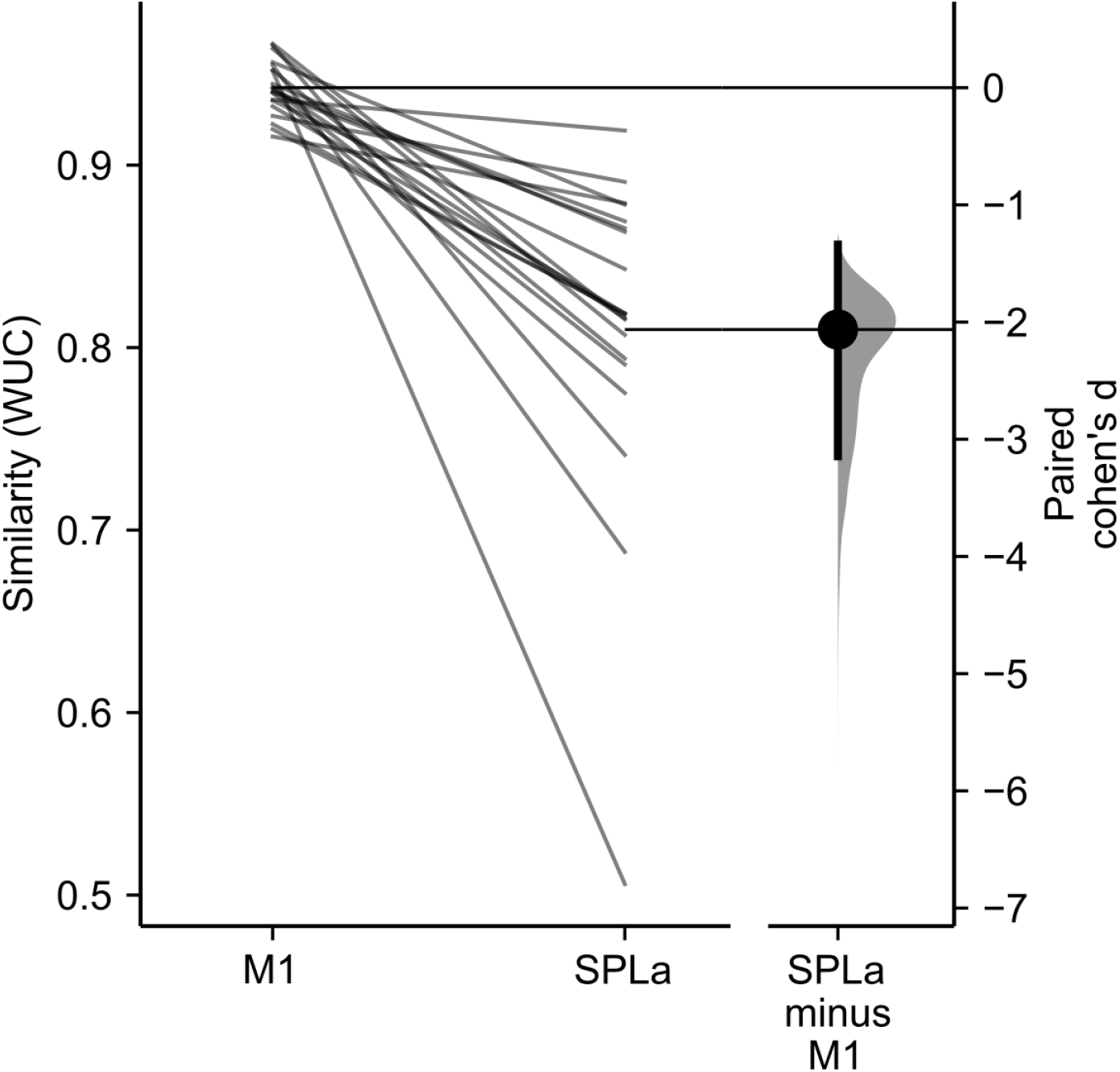
Finger representational structure of tetraplegic individual matches M1 over SPLa for all fMRI participants. Paired Gardner-Altman estimation plot^72^ of the similarity (WUC) between PX (average RDM across sessions) and each fMRI participant. The slopegraph’s connected points (left) show each fMRI participant’s (N = 20) M1/SPLa match with PX’s finger representational structure. Mean difference (right) presented as Cohen’s d (N = 5000 bootstrap samples). Related to Figure 2c and Supplementary Figure 7.

**Supplementary Figure 10.**
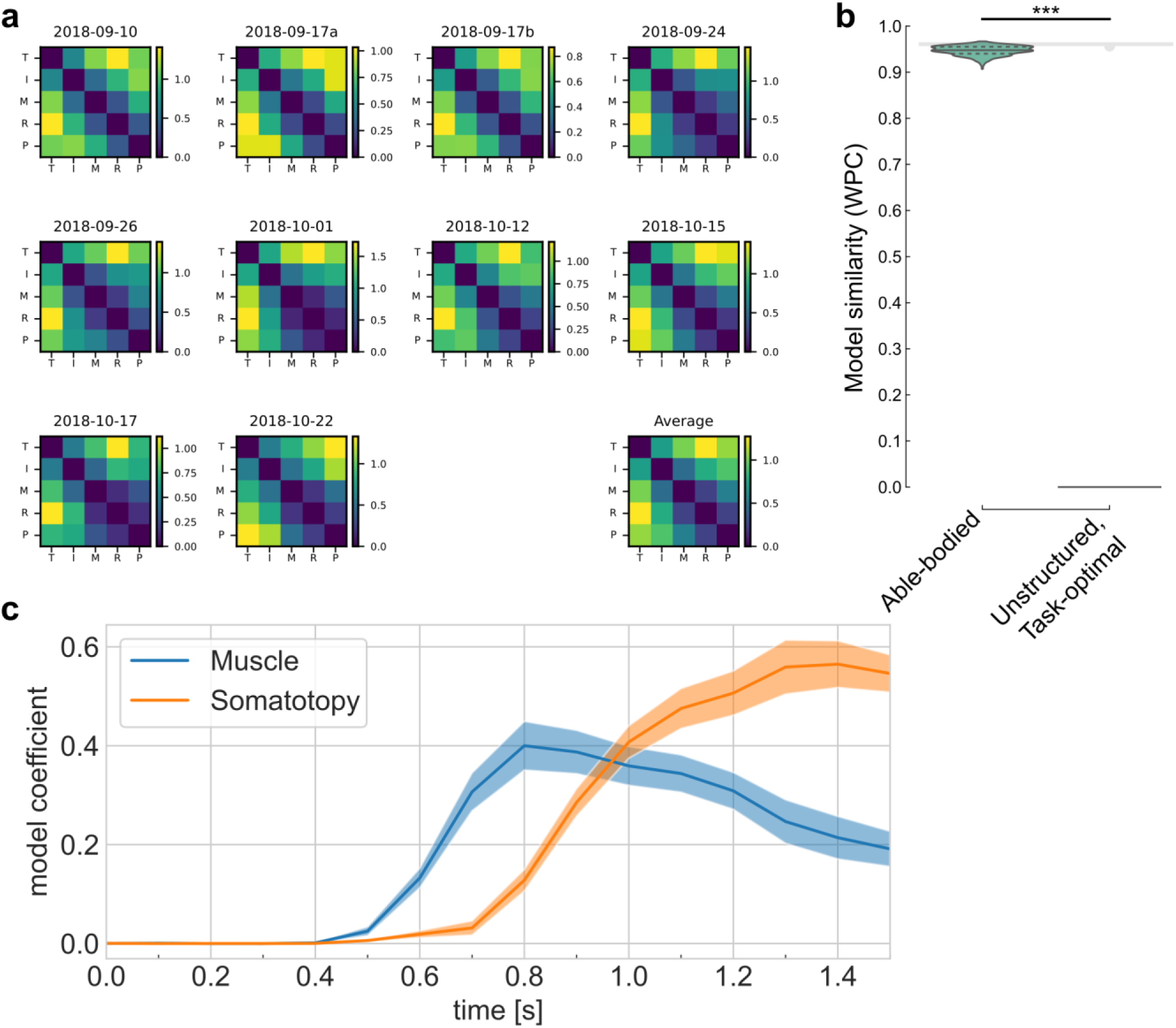
Representational structure during BCI finger control matches the structure of able-bodied individuals when using alternative analysis parameters. (**a**) RDMs calculated with an alternative dissimilarity metric: cross-validated Poisson KL-divergence^63^. Units: nats / neuron. Related to Supplementary Figure 8a and Figure 2e. (**b**) Fit between measured RDMs and motor-intact BOLD data using alternative metrics. Distance metric: cross-validated Poisson KL-divergence. Similarity metric: whitened RDM Pearson correlation^27^. Similar to Figure 2g. Able-bodied model correlation was significantly above the unstructured model (zero correlation) (P = 3.7 × 10^-15^, two-tailed t-test). (**c**) Representational dynamics calculated with an alternative dissimilarity metric: cross-validated Poisson KL-divergence. Similar to Figure 4e.

**Supplementary Figure 11.**
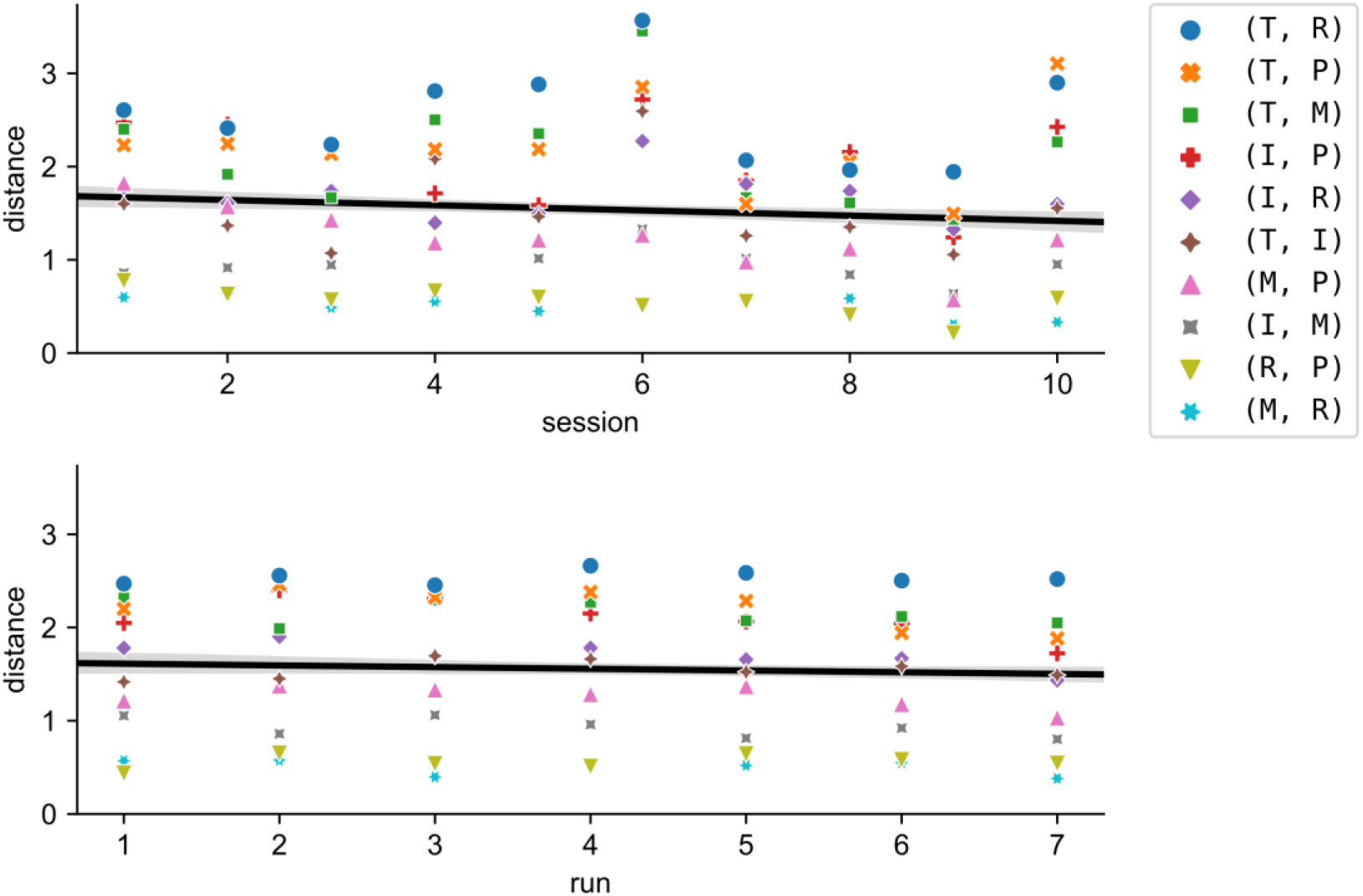
Inter-finger distances did not increase across sessions or within sessions. BCI classification errors might have encouraged inter-finger distances to increase to improve separability, but this did not occur. Inter-finger distances instead decreased slightly (across sessions: t(10) = −4.0, two tailed t-test P < 0.001; within sessions: *t*(10) = −2.4, two-tailed t-test P = 0.017), although the effect size was very small (across sessions: Cohen’s = 0.008; within sessions: = 0.005). Markers indicate average pairwise distance for each finger pair and session (top) or run-within-session (bottom).

**Supplementary Figure 12.**
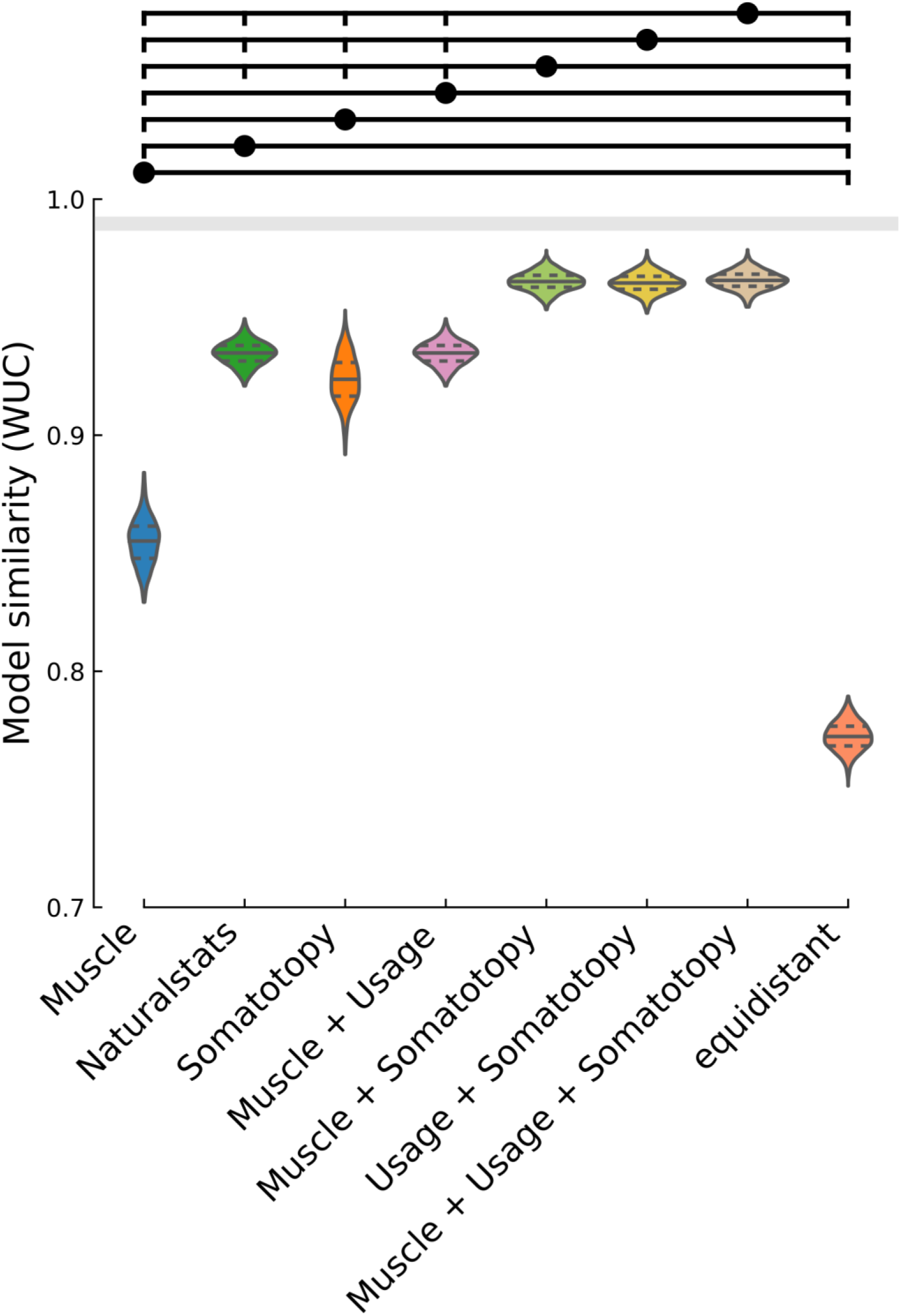
Fit between measured RDM and linear combinations of models. Violin plot of WUC similarity between the measured RDM (N = 1000 bootstrap samples over 10 sessions) and the corresponding model combination. Violin plot: solid horizontal lines indicate the mean WUC over bootstrap samples, and dotted lines indicate the first and third quartiles. Horizontal lines (above) indicate significance groups, where the circle-indicated model is significant over the vertical-tick-indicated models (two-tailed t-test, q < 0.01, FDR-corrected for 28 model-pair comparisons). For example, the muscle+somatotopy combined model is significant over the individual muscle, hand usage, somatotopy, combined muscle+hand-usage, and equidistant (null) models.

**Supplementary Figure 13.**
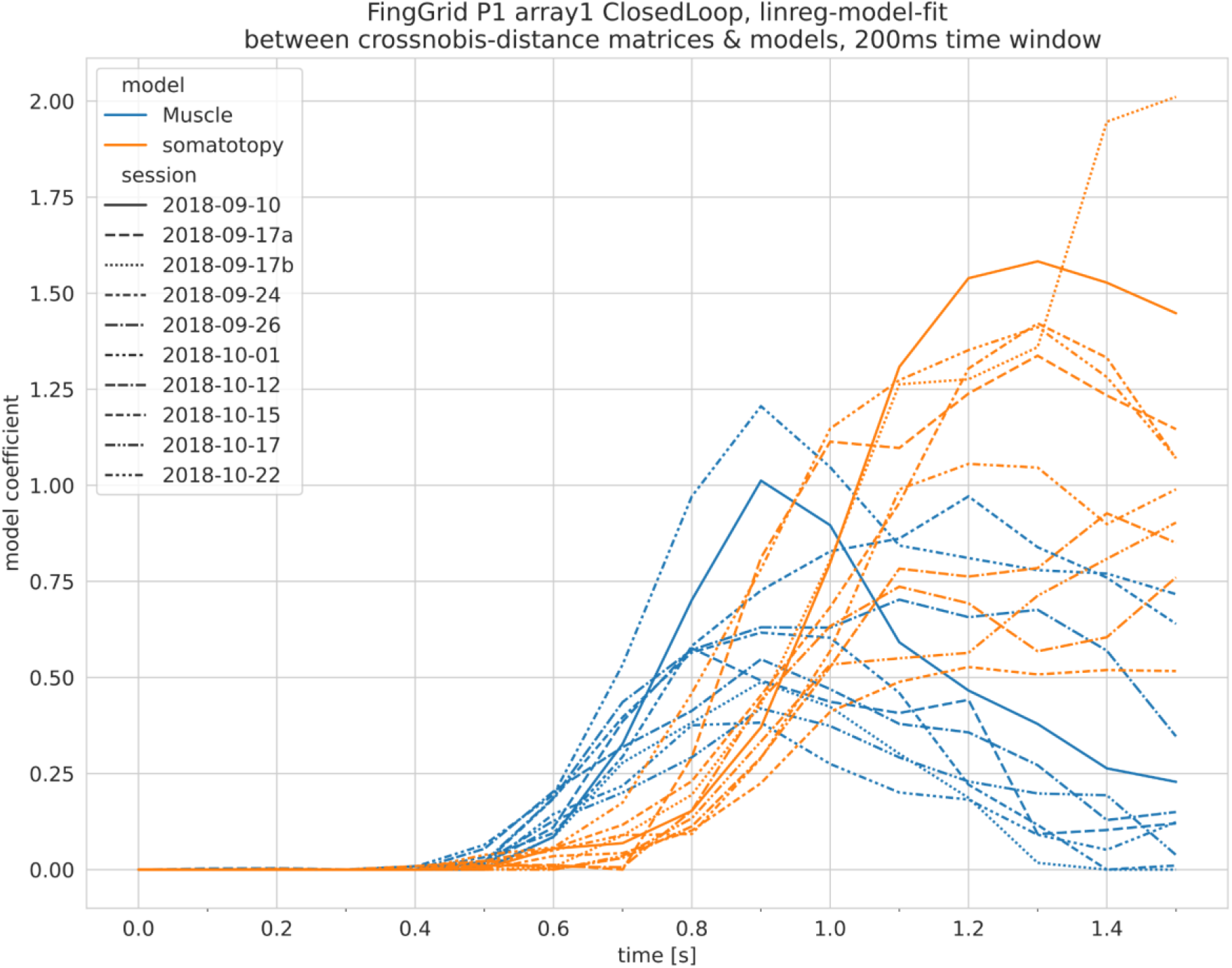
Temporal delays between component models are consistent across single sessions. When linear modeling within single sessions, the muscle model (blue) consistently preceded the somatotopy model (orange). Time difference: 170ms +/- 66ms (s.d. across sessions) (P = 0.002, two-sided Wilcoxon signed-rank test). Line styles indicate session. Related to Figure 4e.

**Supplementary Figure 14.**
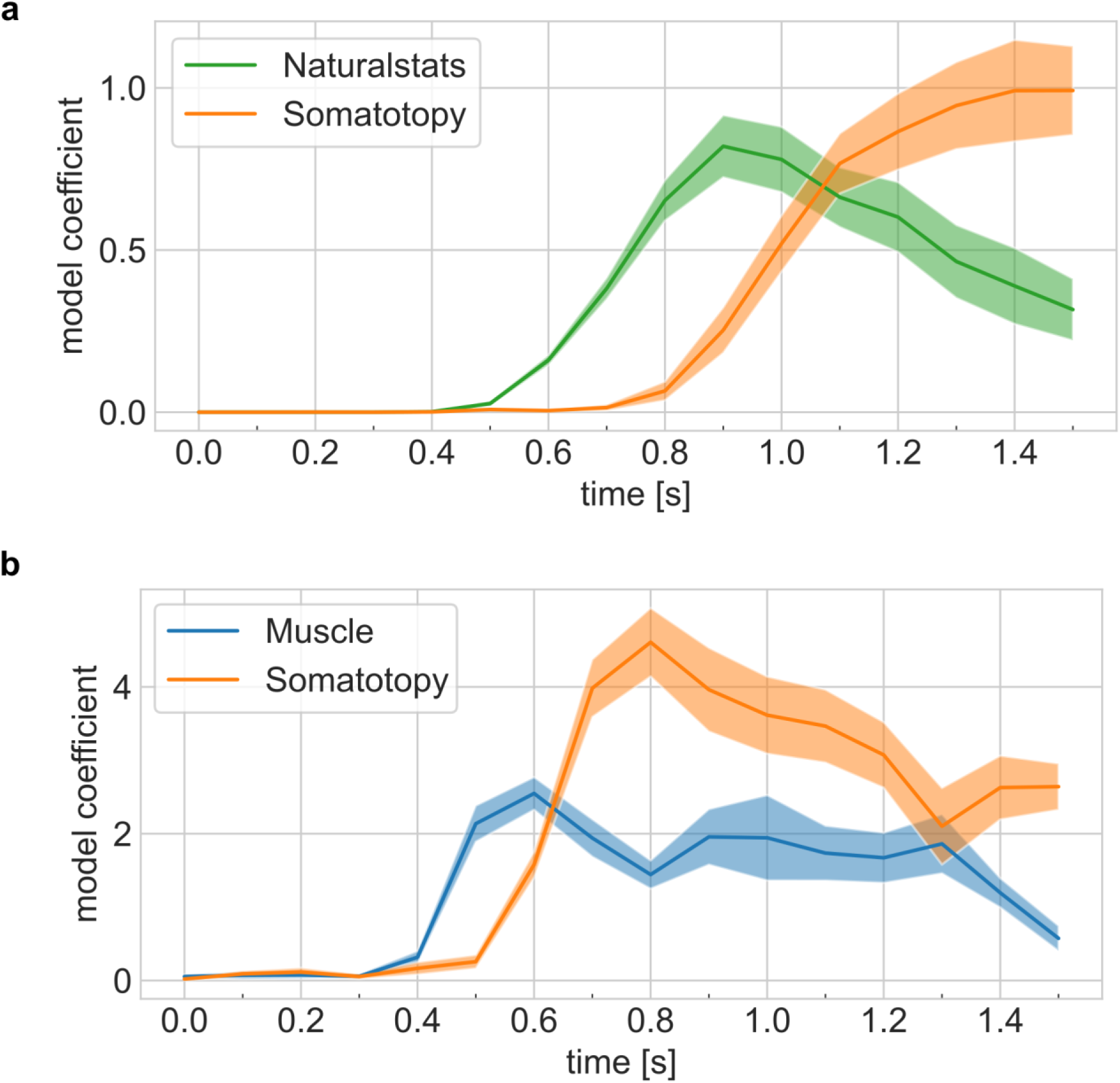
Representational dynamics are robust across tasks and model combination choices. (**a**) Representational dynamics analysis shows an early fit to the hand-usage model and a late fit to the somatotopy model. Confidence intervals indicate +/- s.e.m. across sessions. Related to Figure 4e. (**b**) Representational dynamics analysis shows a consistent delay between models during the calibration task. Note: the absolute timing differs from the main task because the calibration task does not require an initial saccade to read the cue. Related to Figure 4e.

**Supplementary Figure 15.**
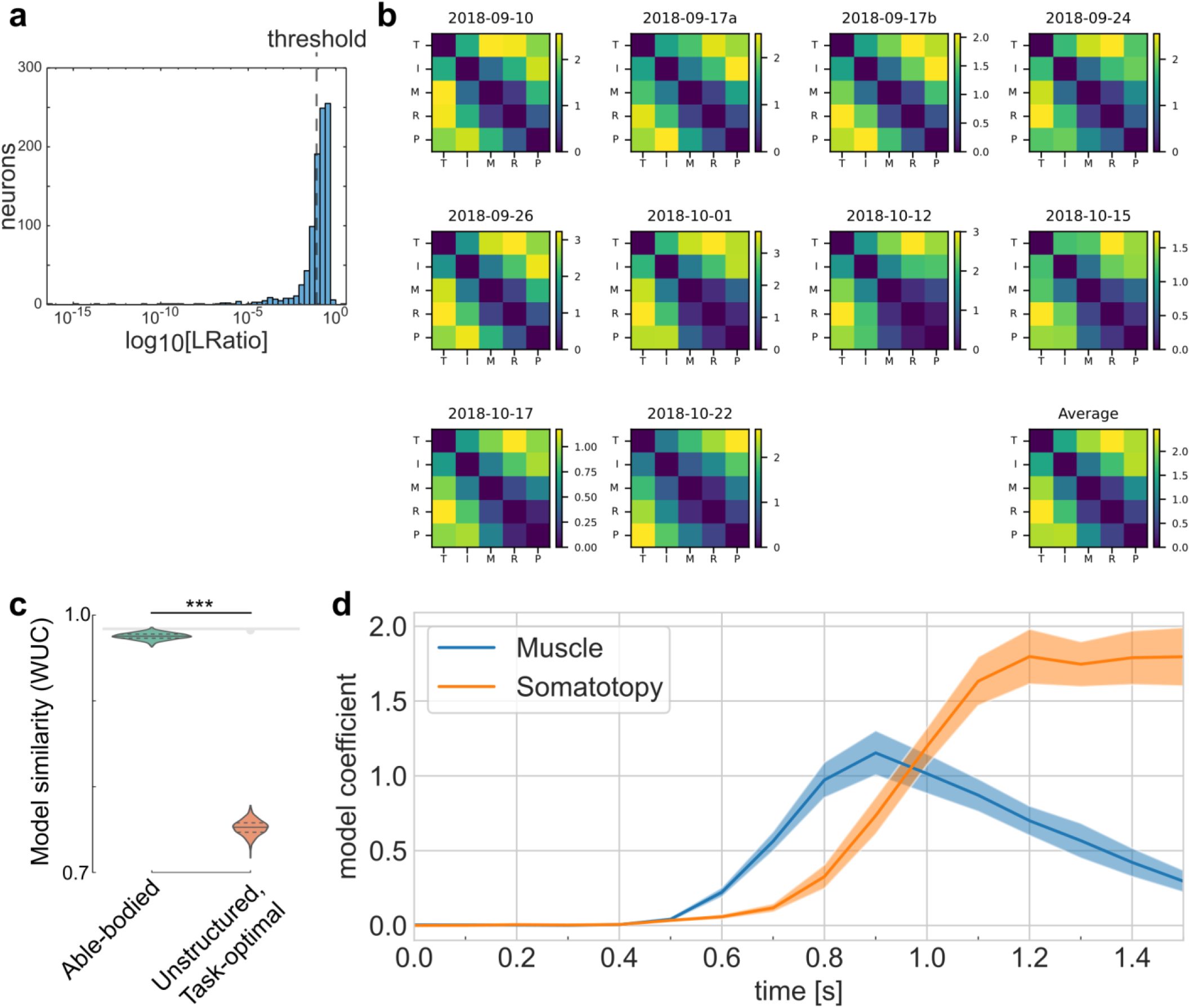
Well-sorted single neurons of the tetraplegic participant match the finger representational structure of able-bodied individuals. (**a**) Histogram of L-ratio, a spike-sorting cluster metric. Threshold for well-isolated units, 33% quantile (L_ratio_ < 10^-1.1^). (**b**) Representational dissimilarity matrices calculated only using well-isolated units, using the cross-validated Mahalanobis distance. Similar to Figure 2e and Supplementary Figure 8a. (**c**) Whitened unbiased similarity (WUC) between measured (B) RDMs (using only well-isolated units) and model predictions (Figure 2b-c), showing that the measured RDMs match the able-bodied BOLD RDM significantly better than they match the unstructured/null model. Error bars indicate +/- s.e.m. Significance indicator: The dot connected to the vertical tick indicates that the able-bodied model fits the data significantly better than the unstructured model (P = 3.1 × 10^-10^, two-tailed t-test). Noise ceiling: Gray region estimates the best possible model fit (Methods). Gray downward-semicircle indicates that the noise ceiling is significantly higher (P < 0.01) than the fit of the unstructured model. Similar to Figure 2g. (**d**) Representational dynamics analysis, repeated using only well-isolated units, shows an early fit to the muscle model and a late fit to the somatotopy model. Confidence intervals indicate +/- s.e.m. across sessions. Similar to Figure 4e.

